# The impact of natural selection on the evolution and function of placentally expressed galectins

**DOI:** 10.1101/505339

**Authors:** Zackery A. Ely, Jiyun M. Moon, Gregory R. Sliwoski, Amandeep K. Sangha, Xing-Xing Shen, Abigail L. Labella, Jens Meiler, John A. Capra, Antonis Rokas

## Abstract

Immunity genes have repeatedly experienced natural selection during mammalian evolution. Galectins are carbohydrate-binding proteins that regulate diverse immune responses, including maternal-fetal immune tolerance in placental pregnancy. Seven human galectins, four conserved across vertebrates and three specific to primates, are involved in placental development. To comprehensively study the molecular evolution of these galectins both across mammals and within humans, we conducted a series of between-and within-species evolutionary analyses. By examining patterns of sequence evolution between species, we found that primate-specific galectins showed uniformly high substitution rates, whereas two of the four other galectins experienced accelerated evolution in primates. By examining human population genomic variation, we found that galectin genes and variants, including variants previously linked to immune diseases, showed signatures of recent positive selection in specific human populations. By examining one nonsynonymous variant in Galectin-8 previously associated with autoimmune diseases, we further discovered that it is tightly linked to three other nonsynonymous variants; surprisingly, the global frequency of this four-variant haplotype is ∼50%. To begin understanding the impact of this major haplotype on Galectin-8 protein structure, we modeled its 3D protein structure and found that it differed substantially from the reference protein structure. These results suggest that placentally expressed galectins experienced both ancient and more recent selection in a lineage-and population-specific manner. Furthermore, our discovery that the major Galectin-8 haplotype is structurally distinct from and more commonly found than the reference haplotype illustrates the significance of understanding the evolutionary processes that sculpted variants associated with human genetic disease.

## Introduction

Galectins are carbohydrate-binding proteins that play diverse and crucial roles in immunity and development (Houzelstein et al. 2004; Rabinovich and Toscano 2009; Than et al. 2015). The immunoregulatory functions of galectins span both innate and adaptive immunity and include—but are not limited to—pattern recognition, T cell-mediated immunosuppression, regulation of acute and chronic inflammation, and selective autophagy in response to pathogen invasion (Rabinovich and Toscano 2009; Vasta 2012). In mammalian development, galectins facilitate implantation and angiogenesis, and they regulate immune tolerance at the maternal-fetal interface (Than et al. 2015).

Of the thirteen galectins found in humans, seven are expressed in the placenta and regulate both immunity and development. All seven placentally-expressed galectins fulfill critical roles in pregnancy, from facilitating implantation events (such as blastocyst attachment to the uterine epithelium) to regulating maternal-fetal immune tolerance through apoptosis of activated T cells (Than et al. 2015). Through this regulation of activated T cells, galectins help protect fetal tissues from attack by maternal immune cells (Than et al. 2012), thereby establishing a healthy environment for fetal development.

The seven placentally-expressed galectins can be divided into two groups on the basis of their phylogenetic conservation and patterns of tissue expression. The first group, which we term the “ancient” group, is comprised of *LGALS1* (which makes the Galectin-1 protein), *LGALS3* (Galectin-3), *LGALS8* (Galectin-8), and *LGALS9* (Galectin-9); all these genes are conserved across vertebrates and expressed in most human tissues, including the placenta (Houzelstein et al. 2004). In contrast, the second group, which we term the “placental cluster” group, is comprised of *LGALS13 (*Galectin-13), *LGALS14* (Galectin-14), and *LGALS16* (Galectin-16); these three genes are only present in primates, formed via gene duplication, are found next to each other in a region of human chromosome 19 (hence the “cluster” term), and appear to be specifically expressed in the placenta (Than et al. 2009).

All proteins in the galectin family mediate their functions through a conserved carbohydrate recognition domain (CRD) comprised of approximately 130 amino acids (Rabinovich and Toscano 2009). CRD structure and organization divide the galectin family into three major types: the prototype, the tandem-repeat type, and the chimera type. The seven placentally-expressed galectins span all three subtypes (Fig. 1A). Prototype galectins (all three in the placental cluster group and Galectin-1) contain a single CRD, whereas tandem-repeat type galectins (Galectin-8 and Galectin-9) contain two CRDs with distinct binding profiles connected by a peptide linker. The chimera type Galectin-3 contains one CRD joined to a collagen-like N-terminal domain. While CRD structure and organization differs among galectin types, all galectin CRDs share eight, mostly invariant residues involved in carbohydrate binding (Fig.1B) (Cummings et al. 2017). CRD topology is well-conserved among galectins, with most family members possessing a CRD comprised of a β-sandwich of five-stranded and six-stranded anti-parallel β-sheets (Than et al. 2015). Despite high structural conservation, the CRD amino acid sequence differs enough among family members to confer distinct carbohydrate-binding preferences (Rabinovich and Toscano 2009).

**Figure 1:**
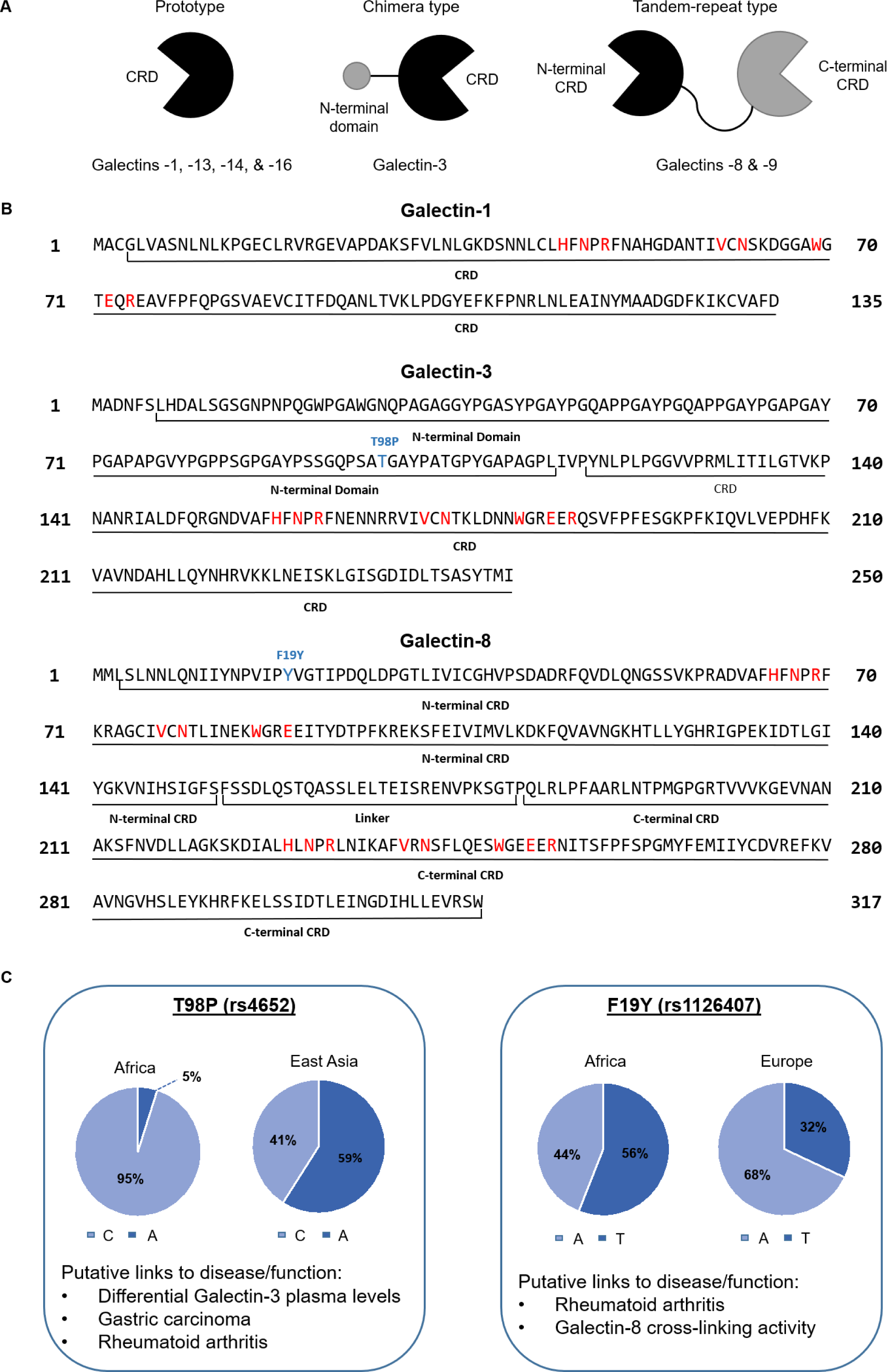
Galectin protein carbohydrate recognition domain (CRD) organization, classification, and sequence. (A) Classification of galectin subtypes based on CRD organization. Prototype galectins are defined by a single CRD, chimera type galectins are defined by a single CRD fused to an additional collagen-like domain, and tandem-repeat type galectins consist of two CRDs joined by a peptide linker of up to 70 amino acids. (B) Amino acid sequences of representatives of each of the three galectin classes. Residues directly involved in carbohydrate binding are highlighted in red and are mostly invariant across all galectin family members. The two residues highlighted in blue represent missense variants described in panel C. (C) Population disparities and putative disease links of the common Galectin-3 single nucleotide variant, rs4652; the C allele, which is at 95% frequency in African genomes, is part of a codon for proline (Pro), whereas the A allele is part of a codon for threonine (Thr). Similarly, the A allele of the common Galectin-8 single nucleotide variant, rs1126407, is part of a codon for tyrosine (Tyr), whereas the T allele is part of a codon for phenylalanine (Phe).

Based on the structural conservation of the galectin CRD and the crucial roles galectins play in fundamental cellular processes, previous studies have hypothesized that mammalian galectins evolved under strong functional constraints that prevented dramatic changes in the carbohydrate recognition domain (Vasta 2012). In contrast, based on the crucial roles galectins play in placental development, other studies have hypothesized that galectins experienced positive selection in early placental mammalian evolution, followed by strong evolutionary conservation thereafter (Than et al. 2008). At present, these hypotheses have been tested via examination of evolutionary rates in only Galectin-1 and the placental cluster galectins (Than et al. 2008, 2009). For example, evidence for positive selection at the origin of placental mammals followed by purifying selection thereafter has been reported for Galectin-1 (Than et al. 2008), but this analysis did not include the other ancient galectins: Galectin-3, Galectin-8, and Galectin-9. Thus, a significant part of placental galectin evolutionary history remains unexamined, and a comprehensive analysis of the evolutionary rates of all placentally expressed galectins could inform us of the tempo and mode of placental galectin evolution during placental mammalian evolution.

Little is known about how galectins evolved *within* the human species. This is surprising given that DNA variants of human immune genes often bear strong signatures of positive selection due to the selective pressures applied by pathogens on the human genome (Barreiro and Quintana-Murci 2010; Fumagalli et al. 2011; Vitti et al. 2013). For example, DARC encodes a membrane protein used by the malaria parasite, *Plasmodium vivax*, for entry into red blood cells (Barreiro and Quintana-Murci 2010). A null mutation in this gene eliminates protein expression and thereby confers full resistance to *P*. *vivax* in homozygotes (Hamblin et al. 2002). Positive selection has targeted the null allele in a population-specific manner, bringing it to fixation in sub-Saharan Africa while it remains virtually absent elsewhere (Hamblin et al. 2002). While natural selection can increase the frequency of such beneficial variants, it can also increase the frequency of disease risk as a byproduct. This phenomenon is well documented for numerous human gene variants that influence susceptibility to autoimmune diseases (Ramos et al. 2015). For example, the *UHRF1BP1* gene experienced positive selection driven by *Mycobacterium tuberculosis*, and it also contains a known risk allele for the autoimmune disease, systemic lupus erythematosus (SLE) (Ramos et al. 2014, 2015). It is thought that this variant influences tuberculosis infection by exerting a regulatory effect on *UHRF1BP1* gene expression (Ramos et al. 2015). Thus, the risk for SLE associated with this variant may have increased as a byproduct of its protection against tuberculosis infection (Raj et al. 2013; Ramos et al. 2015). Human population genetic analyses, such as detection of gene variants highly differentiated between populations, can link disease-associated variants to signatures of natural selection as well as identify tradeoffs (Ramos et al. 2014, 2015), but such approaches have not been used to study placental galectin evolution. This is surprising given that several galectin gene variants have known links to many diseases such as breast cancer and rheumatoid arthritis (Fig. 1C) (Balan et al. 2008; Hu et al. 2011). Examination of the population genetics of galectins in humans could connect these known disease associations to signatures of natural selection and pinpoint new directions for functional studies of galectin gene variants and their role in disease.

To gain insight into the evolution of placental galectins both across mammals as well as in the human population, we performed a comprehensive evolutionary analysis on the seven human galectin genes that regulate both immunity and placental development. We found that members of the ancient group had distinct patterns of nonsynonymous to synonymous substitution rates, suggesting that the functional constraints on galectin evolution were neither uniform across galectin family members nor constant throughout mammalian evolution. By examining patterns of genetic variation in human galectin genes, we identified gene-wide signatures of positive selection on two human galectins. We further found variant-specific signatures of population differentiation, suggestive of geographically-restricted positive selection, for several single nucleotide polymorphisms (SNPs) found in galectin genes.

While studying galectin gene variants, we also observed that the reference sequence commonly used when studying Galectin-8 is not the most common haplotype in the human population. The most common haplotype (hereafter the major haplotype) differs from the current reference sequence by four missense variants. These variants frequently co-occur, and one variant, rs1126407 (hereafter F19Y), is associated with rheumatoid arthritis (Pal et al. 2012). Interestingly, even though F19Y is most often found alongside the three other variants on the major haplotype, it is typically studied on the background of the reference haplotype. To examine differences between proteins encoded by the reference haplotype, major haplotype, and F19Y on the background of reference haplotype, we predicted and compared the protein structures of all three sequences using comparative modeling. The predicted structure of the major haplotype is structurally distinct from that of the reference haplotype as well as from that of F19Y on the background of the reference haplotype. This suggests that the co-occurrence of four variants on the major haplotype leads to structural differences not captured by studies examining F19Y in isolation, highlighting the importance of understanding the genomic background and evolutionary history of loci associated with human complex genetic disease.

## Results

### Ancient Galectin Genes Varied in their Evolutionary Rates During Mammalian Evolution

To examine the evolutionary rates and strength of selection on ancient group galectins, we estimated the ratio of nonsynonymous to synonymous nucleotide substitutions (ω=*d*_N_/*d*_S_) under several different evolutionary hypotheses (Table 1). These hypotheses were designed to test whether galectins experienced accelerated evolutionary rates during the origin of placental mammals (e.g., Hypothesis 4), during the evolution of placental mammals generally (e.g., Hypothesis 3), and/or during the evolution of primates specifically (e.g., Hypotheses 5 and 6). Using the Likelihood Ratio Test, we evaluated the statistical significance of these hypotheses against each other and against a null hypothesis that assumed a uniform ω. Results for hypotheses that differed significantly from the null are summarized in Table 2; all results are detailed in Supplementary Table 1.

**Table 1.** Two sets of evolutionary hypotheses tested for the rate of evolution of placental galectin genes

| Hypotheses for ancient galectins | Parameters |
| --- | --- |
| Null: Uniform rate for all branches | $\omega_{\text{outgroups}} = \omega_{\text{placental stem}} = \omega_{\text{placental clade}} = \omega_{\text{primate clade}}$ |
| 2: Unique rates for placental mammals and placental stem branch | $\omega_{\text{placental stem}} \neq \omega_{\text{outgroups}} \neq \omega_{\text{placental clade}} = \omega_{\text{primate clade}}$ |
| 3: Unique rate for placental mammals | $\omega_{\text{placental stem}} = \omega_{\text{outgroups}} \neq \omega_{\text{placental clade}} = \omega_{\text{primate clade}}$ |
| 4: Unique rate for placental stem branch | $\omega_{\text{placental stem}} \neq \omega_{\text{outgroups}} = \omega_{\text{placental clade}} = \omega_{\text{primate clade}}$ |
| 5: Unique rate for primates | $\omega_{\text{placental stem}} = \omega_{\text{outgroups}} = \omega_{\text{placental clade}} \neq \omega_{\text{primate clade}}$ |
| 6: Unique rates for placental mammals and primates | $\omega_{\text{placental stem}} = \omega_{\text{outgroups}} \neq \omega_{\text{placental clade}} \neq \omega_{\text{primate clade}}$ |
| Hypotheses for placental cluster galectins | Parameters |
| Null: Uniform rate for all branches | $\omega_{\text{primate stem}} = \omega_{\text{outgroups}} = \omega_{\text{Galectin-13}} = \omega_{\text{Galectin-14}} = \omega_{\text{Galectin-16}}$ |
| 2: Unique rate for *LGALS13* | $\omega_{\text{primate stem}} = \omega_{\text{outgroups}} = \omega_{\text{Galectin-14}} = \omega_{\text{Galectin-16}} \neq \omega_{\text{Galectin-13}}$ |
| 3: Unique rate for *LGALS14* | $\omega_{\text{primate stem}} = \omega_{\text{outgroups}} = \omega_{\text{Galectin-13}} = \omega_{\text{Galectin-16}} \neq \omega_{\text{Galectin-14}}$ |
| 4: Unique rate for *LGALS16* | $\omega_{\text{primate stem}} = \omega_{\text{outgroups}} = \omega_{\text{Galectin-13}} = \omega_{\text{Galectin-14}} \neq \omega_{\text{Galectin-16}}$ |
| 5: Unique rates for *LGALS13*, *LGALS14*, and *LGALS16* | $\omega_{\text{primate stem}} = \omega_{\text{outgroups}} \neq \omega_{\text{Galectin-13}} \neq \omega_{\text{Galectin-14}} \neq \omega_{\text{Galectin-16}}$ |
| 6: Unique rate for primate stem branch | $\omega_{\text{primate stem}} \neq \omega_{\text{outgroups}} = \omega_{\text{Galectin-13}} = \omega_{\text{Galectin-14}} = \omega_{\text{Galectin-16}}$ |
| 7: Unique rates for primate stem branch and primate clade | $\omega_{\text{primate stem}} \neq \omega_{\text{outgroups}} \neq \omega_{\text{Galectin-13}} = \omega_{\text{Galectin-14}} = \omega_{\text{Galectin-16}}$ |
| 8: Unique rate for primate clade | $\omega_{\text{primate stem}} = \omega_{\text{outgroups}} \neq \omega_{\text{Galectin-13}} = \omega_{\text{Galectin-14}} = \omega_{\text{Galectin-16}}$ |

**Table 2.** Statistically significant differences in the rate of evolution of placental galectin genes

| Gene | Hypothesis<sup>a</sup> | Likelihood<br>(-lnL) | *p*-value | $\omega$ values | | | |
| --- | --- | --- | --- | --- | --- | --- | --- |
|  |  |  |  | Outgroups | Placental<br>Clade | Placental<br>Stem | Primates |
| ***LGALS1*** | Null | 5353.76 | N/A | 0.15 | 0.15 | 0.15 | 0.15 |
|  | 2 | 5344.95 | 1.5E-04 | 0.20 | 0.16 | 0.01 | 0.16 |
|  | 4 | 5345.53 | 5.0E-05 | 0.17 | 0.17 | 0.01 | 0.17 |
| ***LGALS3*** | Null | 7886.70 | N/A | 0.25 | 0.25 | 0.25 | 0.25 |
|  | 2 | 7834.11 | 1.44E-23 | 0.07 | 0.39 | 0.21 | 0.39 |
|  | 3 | 7834.84 | 2.33E-24 | 0.08 | 0.39 | 0.08 | 0.39 |
|  | 4 | 7884.15 | 2.39E-02 | 0.26 | 0.26 | 0.08 | 0.26 |
|  | 5 | 7873.09 | 1.82E-07 | 0.22 | 0.22 | 0.22 | 0.58 |
|  | 6 | 7831.69 | 1.29E-24 | 0.07 | 0.35 | 0.07 | 0.57 |
| ***LGALS8*** | Null | 8161.84 | N/A | 0.19 | 0.19 | 0.19 | 0.19 |
|  | 5 | 8153.47 | 4.29E-05 | 0.17 | 0.17 | 0.17 | 0.36 |
|  | 6 | 8153.36 | 2.1E-04 | 0.18 | 0.17 | 0.18 | 0.36 |
| ***LGALS9*** | Null | 14016.54 | N/A | 0.34 | 0.34 | 0.34 | 0.34 |
|  | 2 | 14007.70 | 1.4E-04 | 0.28 | 0.36 | 0.11 | 0.36 |
|  | 3 | 14010.44 | 4.74E-04 | 0.24 | 0.36 | 0.24 | 0.36 |
|  | 4 | 14009.79 | 2.36E-04 | 0.35 | 0.35 | 0.11 | 0.35 |
|  | 5 | 14014.56 | 0.0461 | 0.32 | 0.32 | 0.32 | 0.40 |
|  | 6 | 14009.85 | 0.0012 | 0.24 | 0.35 | 0.24 | 0.40 |

<sup>a</sup> Numbers correspond to the hypotheses described in Table 1.

All alternative hypotheses differed significantly from the null hypothesis for *LGALS3* (Table 2; Supplementary Table 1). However, the model assigning a unique ω to the placental stem branch, placental mammals (including primates) and outgroups did not differ significantly from the model in which the placental stem had the same ω as the outgroups (Likelihood Ratio Test or LRT; *p*-value=0.23), so we conclude that Galectin-3 did not experience a different evolutionary rate at the origin of placental mammals. The model assigning a different ω to outgroups, placental mammals (excluding primates), and primates was significantly better than the one assigning different ω to outgroups and placental mammals (including primates) (LRT; *p*-value=0.01), so we conclude that Galectin-3 experienced different evolutionary rates in placental mammals generally and primates specifically. Based on these tests, we conclude that Galectin-3 experienced a fivefold increase in ω in placental mammals (other than primates) relative to outgroups (ω_placental mammals_=0.35; ω_outgroups_=0.07), and experienced a further increase in ω in primates (ω_primates_=0.57) relative to other placental mammals (Fig. 2B).

**Figure 2:**
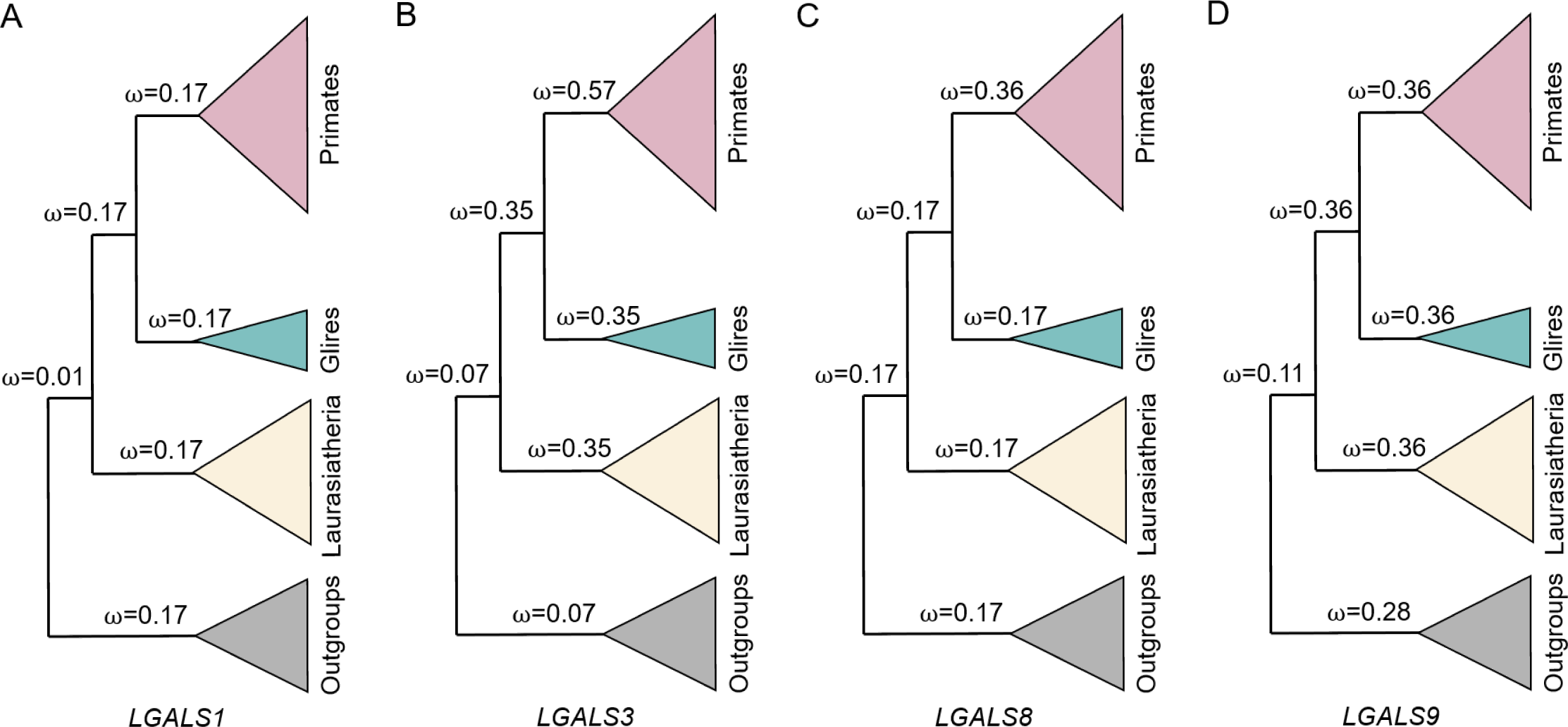
Changes in the evolutionary rate (measured by the ω ratio) in ancient placentally expressed galectin genes. The name of each ancient galectin gene is provided below each topology. The placental stem branch is located at the node representing the split between Laurasiatheria and other placental mammals (Glires and primates). All analyses were performed using codeML (Yang 2007). The complete set of hypotheses tested for each of the ancient galectin genes are provided in Table 1 and their results in Table 2.

Similar to *LGALS3*, all alternative models for *LGALS9* differed significantly from the null hypothesis. The model assigning a unique rate to the placental stem branch, placental mammals (including primates), and outgroups was significantly better than other models assigning any of these variables the same rate, so we conclude that *LGALS9* experienced changes in evolutionary rate at the origin of placental mammals and during the evolution of placental mammals thereafter. Specifically, *LGALS9* experienced an episodic reduction from ω=0.28 to ω=0.11 on the placental stem branch, and then a long-term increase to ω=0.36 during the evolution of placental mammals (Fig. 2D).

In contrast to *LGALS3* and *LGALS9, LGALS8* and *LGALS1* only had two alternative hypotheses that were significantly better than the null hypothesis (Table 2; Supplementary Table 1). For *LGALS8*, we found that the model assigning different ω ratios to both placental mammals (excluding primates) and primates was not significantly better than the model assigning a single ω ratio across primates (LRT; *p*-value=0.64). Thus, we conclude that *LGALS8* experienced accelerated evolution in primates (from ω=0.17 to ω=0.36; Fig. 2C).

For *LGALS1*, we found that the hypothesis assigning different ω ratios to the placental stem, outgroups, and placental mammals (including primates) did not differ significantly from the hypothesis assigning a different ω ratio to only the placental stem (LRT; *p*-value=0.28). Therefore, we conclude that the ω ratio of *LGALS1* was episodically reduced at the origin of placental mammals (from ω=0.17 to ω=0.01; Fig. 2A).

### Placental Cluster Galectins Experienced Similar Evolutionary Rates

To examine the evolutionary rates and strength of selection in placental cluster galectins, we compared the null hypothesis of a uniform ω ratio to alternative hypotheses that posited (1) a different ω for Galectin-13 alone, (2) a different ω for Galectin-14 alone, (3) a different ω for Galectin-16 alone, and (4) a different ω ratio for each of the three genes. None of the alternative hypotheses differed significantly from the null (Supplementary Table 1), suggesting that all three placental cluster galectins evolved at a similar rate of ω=0.60.

Retrieval of placental cluster galectin genes from Ensembl also identified closely related genes named *LGALS13* in pig, *LGALS16* in goat, and *LGALS15* in goat and sheep, as well as additional genes lacking formal names in cow, goat, and opossum (Fig. 3). Our phylogenetic analysis suggests that all these non-primate genes are placed outside the clade formed by placental cluster galectins present in primates (Fig. 3A). Therefore, none of the non-primate genes named *LGALS13* and *LGAS16* are orthologs of the primate *LGALS13* and *LGAS16* genes, respectively. Interestingly, the three goat galectin genes also form a cluster in ∼200 kb region on chromosome 18 and are flanked by genes *EID2* and *DYRK1B*, the same genes used to demarcate and define the cluster of placental galectins found in primates (Than et al. 2009) (Fig. 3B). The same is true for the cow galectins, which also form a cluster and are flanked by the same two genes.

**Figure 3:**
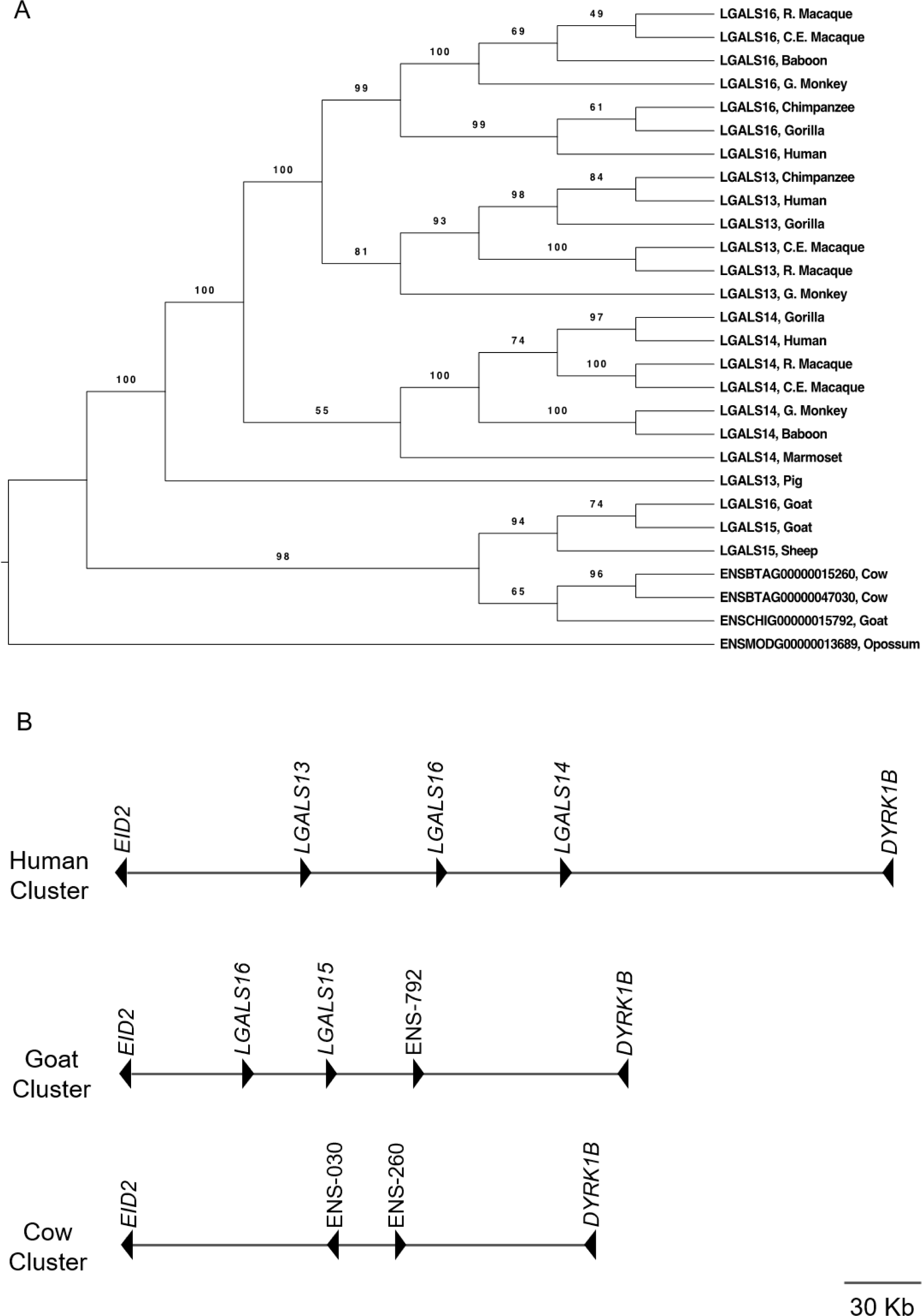
Evolution and genomic organization of placental cluster galectins and related genes. (A) Phylogenetic tree of the placental cluster galectins. Bootstrap support values are included as branch labels. Taxa are labeled as the gene names reported by Ensembl. All alternative models failed to reject the null hypothesis of an equal evolutionary rate (ω=0.60) for the placental cluster galectins. (B) Illustration of genomic organization of galectin gene clusters in human, cow, and goat. Remarkably, even though the cow and goat galectin gene clusters originated independently from the human galectin cluster, all three clusters are flanked by the same two genes: *EID2* on the 5’ end and *DYRK1B* on the 3’ end. Cow and goat genes without official gene names are abbreviated with the last three numbers of their Ensembl accession numbers: ENSBTAG00000015260, ENSBTAG00000047030, and ENSCHIG00000015792.

Therefore, it appears that the placental cluster of galectin genes in primates originated independently of the cluster of galectin genes observed in cow and goat, as has been observed for other clusters of tandemly duplicated genes (Ortiz and Rokas 2017). We tested whether placental cluster galectins specific to primates experienced a different evolutionary rate than these closely related non-primate sequences. We further tested whether the ancestral gene of the placental cluster galectins experienced a short-term change in ω on the primate stem branch. These tests also failed to reject the null hypothesis (Supplementary Table 1), suggesting that the primate-specific galectins evolved at a rate similar to their non-primate relatives.

### LGALS3 and LGALS13 Experienced Recent Positive Selection in Humans

To examine more recent signatures of positive selection that occurred specifically within the human lineage, we calculated three different metrics (Tajima’s *D, F*_ST_, and *H12*) for galectin genes. Tajima’s *D* is optimal for detecting hard selective sweeps that occurred ∼250,000 years ago. Hard sweeps take place when a beneficial variant spreads to very high frequency or fixation alongside the neutral variants linked to it, resulting in the presence of one dominant extended haplotype in the population (Tajima 1989; Sabeti et al. 2006). The fixation index *F*_ST_ is optimal for detecting population differentiation, which can result from local adaptation, that occurred 50,000 – 75,000 years ago (Weir and Cockerham 1984). Finally, the *H12* index is optimal for detecting soft selective sweeps, which take place when multiple beneficial variants of the same locus spread into the population, resulting in two or more dominant extended haplotypes increasing in frequency in the population (Pennings and Hermission 2006; Messer and Petrov 2013; Garud et al. 2015).

To assess statistical significance, we simulated sequences of similar length to galectins based on a neutral evolution model that considered human population demography (see Methods). We specifically simulated the genotypes of 661, 503, and 504 individuals to reflect the number of individuals analyzed from the African, European, and East Asian populations, respectively, in the 1000 Genomes Project. Analysis of the simulated sequences enabled us to generate null distributions of values for each metric across each simulated galectin gene. We then compared the galectin genes’ actual values against the null distributions to evaluate significance. We found that *LGALS3* experienced positive selection in the form of a soft selective sweep (*H12*=0.12; empirical *p*-value<0.01), while *LGALS13* experienced a hard selective sweep (Tajima’s *D*=-2.32; empirical *p*-value=0.01) (Fig. 4A & B). The other galectin genes, such as *LGALS1* (Fig. 4C & D), did not show signatures of positive selection for any of the three metrics, suggesting that they evolved neutrally in the human lineage (Supplementary Table 2).

**Figure 4:**
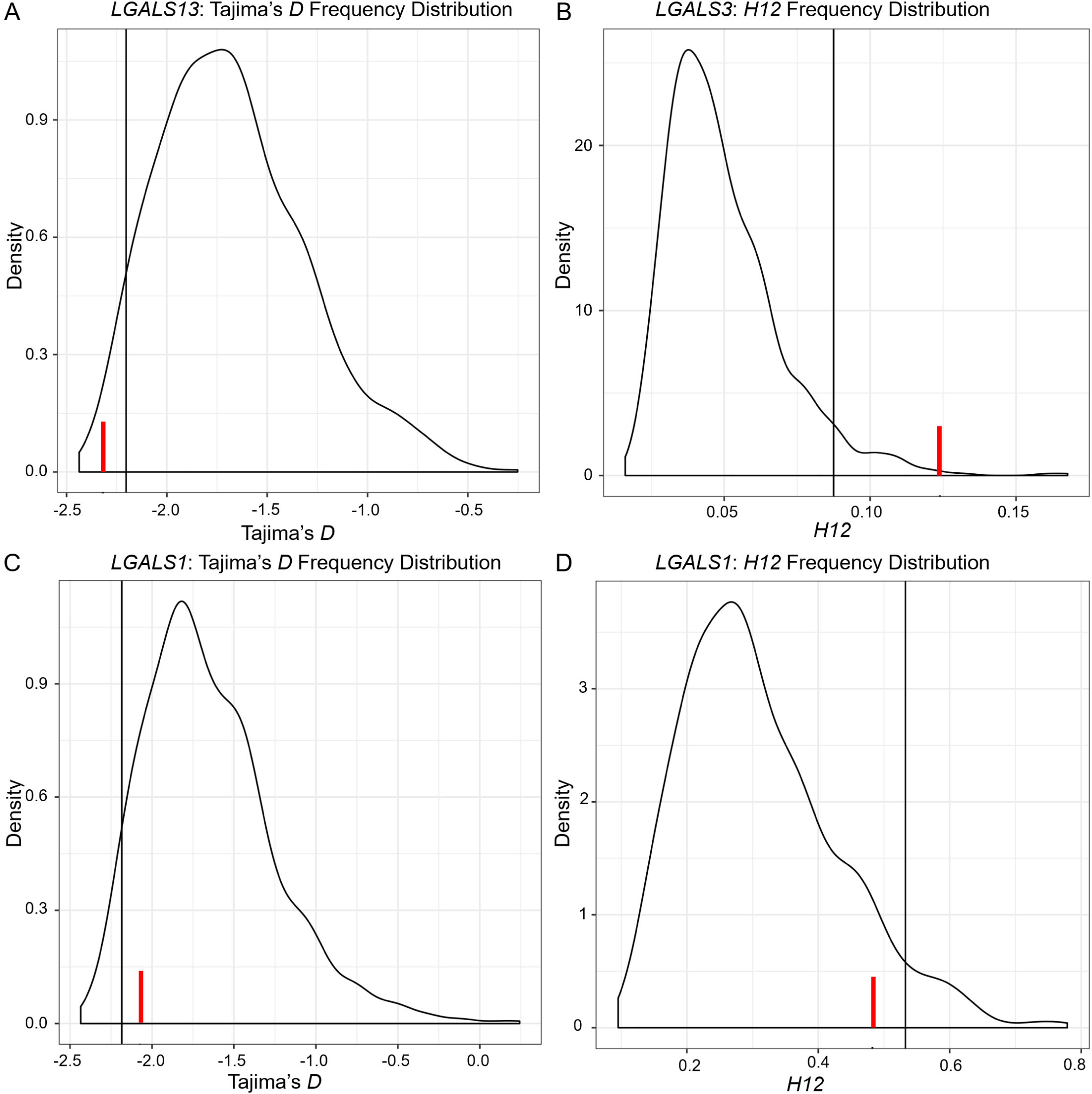
Representative distributions of Tajima’s *D* and *H12*. Vertical lines indicate the significance cutoff corresponding to the 95^th^ percentile of the null distribution. Red bars indicate where the actual values calculated for galectin genes fall along the distribution. Significantly negative values of Tajima’s *D* are interpreted as evidence of positive selection via hard selective sweeps, whereas significantly positive values of *H12* are interpreted as evidence of positive selection via soft selective sweeps. (A) A significantly negative value of Tajima’s *D* is found for *LGALS13*. (B) A significantly positive value of H12 is found for *LGALS3*. (C) and (D) *LGALS1* did not show significant results for these statistics.

### Several Galectin SNPs Previously Linked to Disease Show Extreme Population Differentiation

Calculating *F*_ST_ for specific SNPs can identify signals of positive selection (Barreiro & Quintana-Murci 2010) that may be masked in gene-wide analyses due to the presence of other, neutrally evolving SNPs in a gene. To detect SNP-specific extremes in *F*_ST_, we compared galectin SNP *F*_ST_ values to neutral SNP *F*_ST_ distributions based on the same population comparison (e.g., a galectin SNP *F*_ST_ value calculated by comparing East Asians and Africans would be evaluated against an *F*_ST_ distribution for SNPs from neutral sequences simulated for East Asia and Africa). We found that 138 galectin SNPs exceeded the 95^th^ percentile of simulated *F*_ST_ for at least one population comparison (Supplementary Table 3). Since the functional consequences of the majority of these SNPs are unknown, we further filtered the SNPs to include only those seven that had a putative link to a disease or phenotype, and we estimated their minimum allele ages using parsimony ancestral reconstruction across a phylogenetic tree of 30 mammals (see Methods for detail). Finally, we submitted missense variants to PolyPhen-2 (Adzhubei et al. 2010) for preliminary variant effect prediction. The *F*_ST_ values, PolyPhen values, and allele ages of these SNPs are described in Table 3. We found seven galectin SNPs that exhibit significant population differentiation (as well as two additional SNPs that are marginally non-significant) and that are linked to disease or another phenotype.

**Table 3.** *F*_ST_ values, phenotype links, and allele ages of placental galectin variants with significant population differentiation and putative links to disease or other phenotypes

| SNP | Gene | $p$-value | Functional consequence | Highest calculated $F_{ST}$ | PolyPhen score | Putative disease/phenotype link | ID and age of oldest allele<sup>c</sup> | ID and age of youngest allele |
| --- | --- | --- | --- | --- | --- | --- | --- | --- |
| **rs4652** | *LGALS3* | 0.004 | Missense (T > P) | 0.52 | 0 (benign) | Several | C; 105 | A; 15.8 |
| **rs10403702** | *LGALS16* | 0.03 | Missense (L > P) | 0.20 | 0 | Acute insulin response to glucose | C; 43 | T; human-specific allele |
| **rs4830** | *LGALS14* | 0.02 | Missense (C > S/R) | 0.32 | 0.02 | Epistatic effect in breast cancer | A; human-specific | T; human-specific |
| **rs1009977** | *LGALS3* | 0.01 | Intron variant | 0.35 | N/A | Cognitive function in old age | G; 29.4 | T; 15.8 |
| **rs10498474** | *LGALS3* | 0.02 | Intron variant | 0.24 | N/A | Body weight | A; 43 | G; human-specific |
| **rs2273865** | *LGALS8* | 0.048 | Nonsense variant | 0.14 | N/A | Differential expression of galectin-8 isoforms | T; 105 | A; human-specific |
| **rs4659682** | *LGALS8* | 0.01 | Intron variant | 0.34 | N/A | ADHD-IV impulsivity | A; 105 | G; human-specific |
| **rs4644<sup>a</sup>** | *LGALS3* | 0.056 | Missense (P > H) | 0.11 | 1 (deleterious) | Several | C; 105 | A; human-specific |
| **rs1126407<sup>a</sup>** | *LGALS8* | 0.066 | Missense (F > Y) | 0.10 | 0.01 | Rheumatoid arthritis | A; 105 | T; 6.23 |

<sup>a</sup> These SNPs are marginally non-significant but are linked to disease, so we opted to include them. <sup>c</sup> All ages are given in millions of years; human-specific alleles are identified as such.

Of these, rs4652, a *LGALS3* variant causing a proline to threonine substitution, showed the highest *F*_ST_ value: 0.52 between African and East Asian populations. It is associated with rheumatoid arthritis (Hu et al. 2011), risk for gastric carcinoma (Shi et al. 2017), differential *LGALS3* gene expression, and differential Galectin-3 circulation levels in the blood (de Boer et al. 2012). We found that both rs4652 alleles are ancient: The C allele is found across eutherian mammals and is estimated to be approximately 105 million years old, whereas the A allele is found across the great apes and is estimated to be approximately 16 million years old. The rs4652 locus has thus been polymorphic for millions of years. This provides interesting evolutionary context for the SNP’s population disparity, in which the C allele has nearly reached fixation (95% frequency) in African populations but remains at less than 50% frequency in non-African populations. This suggests that the ancient C allele has been targeted by selection in Africa, and that the selective force operating there has been powerful enough to overcome the evolutionary forces that maintained the polymorphism for millions of years.

### LGALS8 has Two Major Haplotypes Defined by Four Common Variants

Another interesting variant, rs1126407 (F19Y), causes a phenylalanine to tyrosine substitution in Galectin-8. This variant narrowly missed the threshold for significant population differentiation (*F*_ST_ value=0.1; *p*-value=0.07). The A allele, which encodes tyrosine, has existed for approximately 105 million years, while the younger T allele, which encodes phenylalanine, has existed for approximately 6 million years. The A allele is present at 68% frequency in European populations, while it remains below 50% frequency in Africa (44%) and East Asia (48%). F19Y was associated with rheumatoid arthritis in a previous study (Pal et al. 2012). Two functional studies that followed the association study elucidated both structural and functional impacts of this variant on Galectin-8 carbohydrate-binding activity (Ruiz et al. 2014; Zhang et al. 2015). Importantly, both studies focused on the F19Y variant alone by mutating only F19 to Y19 on a background of the human reference sequence; however, our analysis revealed another variant, R36C (rs1041935), that is both highly differentiated and in complete linkage disequilibrium (D’=1; r^2^=1) with F19Y. Further investigation of 1,000 Genomes Project data revealed that Y19 frequently co-occurs with three other missense variants (C36, V56, and S184) in over 80% of Galectin-8 protein haplotypes. We also found a global frequency of 31.7% for the reference haplotype (F19, R36, M56, and R184) of the canonical isoform typically used in Galectin-8 functional studies; in contrast, the global frequency of the major haplotype (Y19, C36, V56, S184) is 49.6%. Notably, Y19 has never been observed in the reference background (i.e., with R36, M56, and R184) (Fig. 5A), suggesting that the “hybrid” haplotype (Y19, R36, M56, and R184) examined by the studies of Ruiz *et al*. 2014 and Zhang *et al*. 2015 either doesn’t exist in the human population or is present at very low frequency.

**Figure 5:**
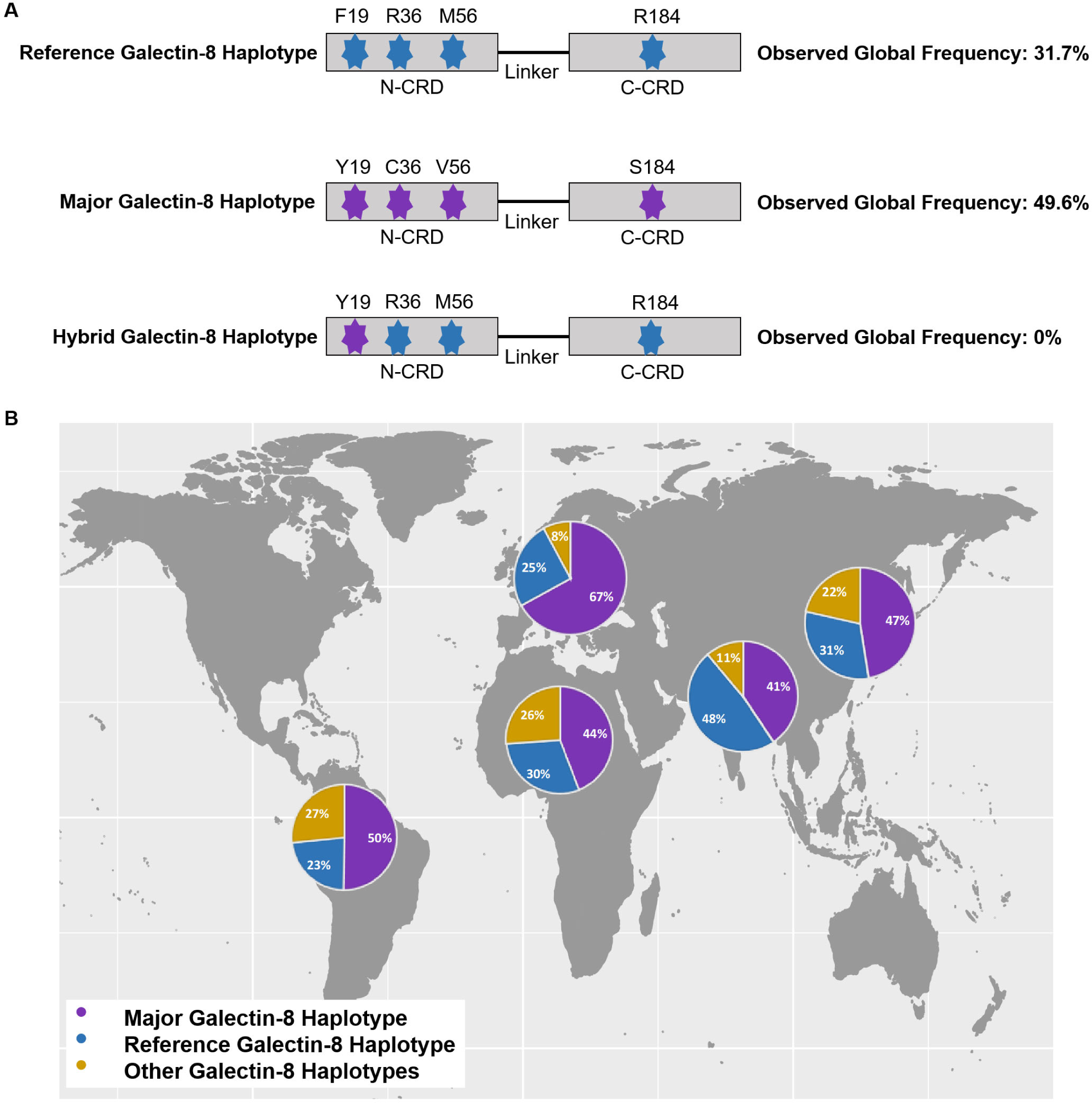
Protein haplotype frequencies of the canonical Galectin-8 isoform across and within five major human populations. (A) Global frequency of two common Galectin-8 protein haplotypes defined by four missense variants. The reference haplotype commonly used in Galectin-8 functional studies is not the most common protein haplotype. None of the 2,504 human genomes in the 1000 Genomes Project contain the hybrid haplotype. (B) Population frequencies of different Galectin-8 protein haplotypes mapped to five major continental populations defined in the 1000 Genomes Project: Africa, America (admixed populations), East Asia, Europe, and South Asia. Frequencies are based on data from the 1000 Genomes Project compiled in Ensembl for the canonical isoform. The Ensembl accession number of the corresponding *LGALS8* transcript is ENST00000341872.10.

### Galectin-8 Sequence Differences Influence Structure

To evaluate potential structural differences between the major, reference, and hybrid haplotypes, we used comparative modeling with RosettaCM to predict their structures (Methods). We compared the models of the three haplotypes using global root-mean-square deviation (RMSD), a metric that quantifies the mean distance between atoms of superimposed protein structures. Our comparisons revealed structural differences between the reference and major haplotypes (RMSD=0.79), between the reference and hybrid haplotypes (RMSD=0.70), and between the hybrid and major haplotypes (RMSD=0.74) (Fig. 6A, B, & C respectively).

**Figure 6:**
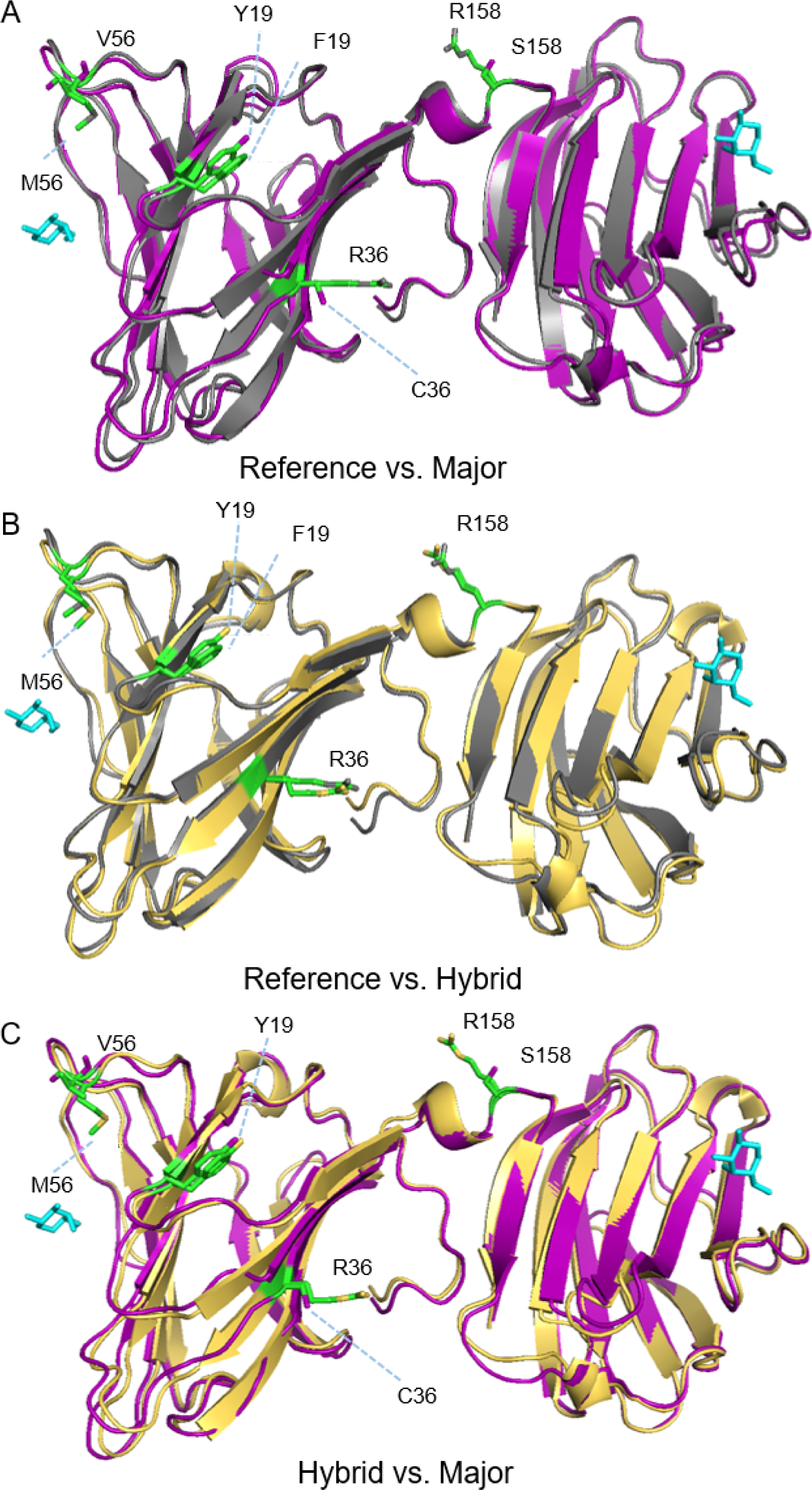
Protein structural differences between the central models of the major haplotype, the reference haplotype, and the hybrid haplotype. The protein structure for the reference haplotype is colored gray, the major haplotype purple, and the hybrid haplotype yellow, respectively. Variant residues are colored green. Cyan-colored structures illustrate the approximate position typically occupied by carbohydrate ligands that bind the carbohydrate recognition domain. RMSD values are given in units of angstroms. (A) Model based on the major haplotype superimposed with the model based on the reference haplotype (RMSD=0.79). (B) Model based on the hybrid haplotype superimposed with the model based on the reference haplotype (RMSD=0.70). (C) Model based on the hybrid haplotype superimposed with model based on the major haplotype (RMSD=0.74).

These results suggest that the major haplotype (Y19, C36, V56, S184) induces different structural changes than the hybrid haplotype (Y19, R36, M56, and R184). However, it is important to note that considerable structural variation exists in the top 10 scoring models of all three sequences (Supplementary Figure 1). RosettaCM is designed to predict the most likely conformations for a protein of interest. However, it is possible that a single model cannot accurately capture the conformational flexibility of Galectin-8. To address this issue, we employed the Calibur clustering algorithm to examine whether the carbohydrate binding region of the top 100 scoring models of each haplotype (i.e., the reference haplotype, the major haplotype, and the hybrid haplotype) group into different structure-based clusters. This would indicate that, despite the variation *within* each set, the different haplotypes are still likely to adopt different structural conformations in the natural protein at the major carbohydrate binding site.

Focusing on residues spanning the binding region of the N-terminal CRD, models based on the hybrid haplotype tend to cluster with models based on the reference haplotype, which together comprise roughly equal amounts of 86% of the models present in the largest cluster (Supplementary Figure 2A). In contrast, models based on the major haplotype tend to cluster on their own, comprising 70% of the models in the second-largest cluster. This reinforces the hypothesis that the major haplotype has different structural impacts than the hybrid haplotype, and it also suggests that these differences are pronounced at the N-terminal CRD (Supplementary Figure 2B), one of two major functional sites for Galectin-8’s carbohydrate-binding activities.

## Discussion

We found that seven placentally expressed galectin genes involved in immunity evolved at different rates during the evolution of placental mammals. *LGALS3, LGALS8*, and *LGALS9* experienced periods of accelerated evolution that reflect changes in functional constraints possibly driven by positive selection. We found further evidence for selection in the human lineage, including gene-wide signatures of positive selection acting on *LGALS3* and *LGALS13*, as well as SNP-wise signatures of population differentiation. By combining these signatures with findings from previous studies, we identified seven galectin gene variants that experienced positive selection and that are linked to disease. Taken together, these results suggest that the evolutionary history of placentally expressed galectin genes has been dynamic, both across mammals as well as within humans.

Prior studies have hypothesized strong evolutionary constraints on the basis of galectins’ fundamental and evolutionary ancient immune functions (Vasta 2012). In contrast, our results show that galectins evolved at different evolutionary rates and experienced both episodic and long-term rate changes, suggesting that functional constraint varied across genes, lineages, and time windows. In the case of *LGALS3, LGALS8*, and *LGALS9*, periods of accelerated evolution are suggestive of relaxed functional constraints. This is especially interesting considering the multifunctional nature of mammalian galectins, which have fundamental immunoregulatory functions that likely predate the origin of placental mammals, but which also play important roles in placental development (Vasta 2012; Than et al. 2015). Periods of increased ω ratios may reflect evolutionary modifications that occurred as these ancient immunity genes adapted their immunity functions for new roles in placental development.

Phylogenetic evidence also suggests that *LGALS1* and *LGALS9* experienced increased purifying selection (decreased ω) at the origin of placental mammals. This result is in contrast to previous findings by Than *et al*. (2008), who found evidence for positive selection acting on *LGALS1* at the origin of placental mammals. This contradiction is explained by differences in methodology: Than *et al*. inferred site-specific positive selection through the use of branch-site models, whereas we inferred negative selection through the use of branch models (Yang 2007). Branch models can miss positive selection that acted on only a subset of codons; branch-site models would not, but have been shown to suffer from high false-positive rates (Pond et al. 2011). By utilizing only branch models, we avoided the risk of false-positives and detected negative selection acting on *LGALS1* and *LGALS9* in the genes at-large.

In contrast to the variable evolutionary rates found in ancient galectin genes, we found evidence for a constant evolutionary rate in placental cluster galectin genes (Figs. 2, 3). Previously established evolutionary and functional evidence may help explain this difference between the ancient and placental cluster galectins. For example, all three of the placental cluster galectins are solely expressed in the placenta (Than et al. 2009), and they evolved relatively recently by tandem duplication in the primate lineage and share specific carbohydrate binding preferences and apoptotic functions (Than et al. 2009). Thus, shared tissue environment and biological function could have produced a similar selection pressure for all three placental galectins, leading to their similar rates of evolution observed in our analysis.

The finding that placental cluster galectins exhibit highly similar evolutionary rates contradicts previous findings from Than *et al*. (2009), who proposed highly variable evolutionary rates for these placental galectins. This inference was based on the free-ratio model that assigns a unique ω to every branch in the phylogeny. This test has received criticism because it is parameter-rich and prone to overfitting; its use is also discouraged in the PAML manual (Yang 2007). Than *et al*. also performed their analysis on a phylogeny containing several paralogs of *LGALS13, LGALS14*, and *LGALS16* that exist in expanded gene clusters in other primate species. By contrast, our analysis only focused on orthologs of the three genes of this cluster that are present in humans and solely expressed in the placenta. Thus, Than *et al*.’s inference of variable evolutionary rates applies to a larger set of genes than the one included in our analysis; however, inferences based on a free-ratio model are still parameter-rich and unlikely to produce accurate estimates of all ω values (Yang 2007), and they are partly contradicted by the failure of branch model tests to reject the null hypothesis in our analysis.

Our analysis widens the lens on the evolution of the placental cluster galectins. By including closely related non-primate genes, we discovered that galectin gene clustering is not unique to primates: both goat and cow contain small clusters of galectin genes in a chromosomal region homologous to the region containing primate placental galectins. Interestingly, *LGALS*15, a galectin gene present in both sheep and goat, is solely expressed in the uterus and supports trophoblast attachment in these species (Lewis et al. 2007). This highlights a striking case of convergent evolution of a cluster of tandem duplicated genes (Ortiz and Rokas 2017) between artiodactyls and primates: both lineages evolved gene clusters of variable size comprised of galectins with unique expression in reproductive tissues and putative functions in pregnancy.

Human immune genes often show strong signatures of positive selection as our genomes evolved in response to selective pressures applied by pathogens (Barreiro and Quintana-Murci 2010). We found that *LGALS3* and *LGALS13* exhibit signatures of recent positive selection in the human lineage. *LGALS3* specifically experienced a soft selective sweep, in which multiple beneficial alleles rise to relatively high frequencies in a population (Pennings and Hermission 2006; Messer and Petrov 2013; Garud et al. 2015). *LGALS3* encodes a highly multifunctional protein with nearly ubiquitous tissue expression (Dumic et al. 2006), and it is highly likely that the selected variants may have impacted several Galectin-3-related processes at once; consequently, identifying which function might have been the driving force behind selection is challenging. It is notable, though, that *LGALS3* also showed accelerated evolution in placental mammals and primates. Taken together, these results suggest that Galectin-3 has long been a substrate for evolutionary modifications, undergoing accelerated substitution rates in ancient mammals and significant changes in haplotype frequencies in modern humans.

We also discovered SNP-specific signatures of positive selection through analysis of the *F*_ST_ statistic, which can detect geographically restricted positive selection (Barreiro and Quintana-Murci 2010). Seven of these SNPs are putatively linked to diseases, such as breast cancer and rheumatoid arthritis, as well as other phenotypes, such as ADHD impulsivity (Kim et al. 2012).

Of these, rs4652, a *LGALS3* variant, showed the strongest signature of population differentiation (*F*_ST_=0.52 between African and East Asian populations). rs4652 has been implicated in diseases such as rheumatoid arthritis and gastric carcinoma (Hu et al. 2011; Shi et al. 2017). After finding that Galectin-3 deletion in mice conferred partial protection to malaria, Oakley *et al*. 2009 hypothesized that this SNP could influence malaria resistance by disrupting Galectin-3 multimerization, an idea strengthened by the fact that rs4652 disrupts a key repeat in the domain required for Galectin-3 multimerization. In addition to influencing Galectin-3’s multimerization, rs4652 may also affect its gene expression and circulation levels. de Boer *et al*. (2012) reported that rs4652 is strongly associated with Galectin-3 circulating levels in the blood. Galectin-3 circulating levels are implicated in several diseases that span immunological diseases and cardiovascular disorders such as arteriosclerosis (de Boer et al. 2012; Madrigal-Matute et al. 2014). Interestingly, Banfer *et al*. (2018) recently elucidated a mechanism for Galectin-3’s exosomal secretion that involves direct binding to Tsg101, a component of the endosomal sorting complex required for transport (ESCRT). Galectin-3 interacts with Tsg101 via a P(S/T)AP peptide motif that is disrupted by rs4652, and a coimmunoprecipitation assay confirmed that the variant altered Galectin-3’s capacity to precipitate Tsg101 (Banfer *et al*. 2018). We propose that this impact may represent a functional link to the variant’s association with Galectin-3 circulation levels in the blood.

Another variant, rs1126407 (F19Y), lies within *LGALS8* and narrowly missed the significance threshold for population differentiation (*p*-value=0.07). This variant is associated with rheumatoid arthritis and has been the subject of several functional analyses (Ruiz et al. 2014; Zhang et al. 2015). These studies analyzed F19Y in the background of the reference human haplotype, which resulted in the exclusion of three other nonsynonymous variants that are linked with it in over 80% of Galectin-8 haplotypes in the human population. This exclusion means that the functional effects of the F19Y variant were examined in the context of a “hybrid” haplotype that is not found in the 5,008 human haplotypes present in the 1,000 Genomes Project. Our results predict that the major haplotype, containing all four variants, impacts the protein structure differently than the hybrid haplotype, and they predict that the major haplotype induces a greater impact on the key binding region of Galectin-8’s N-terminal carbohydrate recognition domain (CRD). This could thus impact Galectin-8’s binding affinity with a variety of ligands that bind its N-terminal CRD. These results illustrate the importance of considering human haplotype diversity when investigating natural protein function and disease-associated genetic variants. By referring only to the reference sequence, investigators may not capture the properties of far more common, non-reference sequences that may harbor several amino acid differences in the protein of interest.

Future studies of Galectin-8 sequence and structure must consider its haplotype diversity, in particular the major haplotype, which is more common globally than the reference sequence. Future studies of the structure of Galectin-8 in complex with the glycoprotein CD44 would also be valuable. Galectin-8 normally executes an anti-inflammatory function by inducing apoptosis of synovial fluid (SF) cells through binding with CD44 at the cell surface (Sebban et al. 2007). In rheumatoid arthritis, SF cells express a unique CD44 variant (CD44vRA) that has been shown to ligate and sequester Galectin-8 in patients with rheumatoid arthritis (Sebban et al. 2007), thereby reducing its anti-inflammatory effects. We hypothesize that major Galectin-8 haplotypes have differential affinity for binding CD44vRA, and that this explains the association of F19Y with rheumatoid arthritis.

## Materials and Methods

### Ortholog Retrieval and Sequence Alignment

We examined seven human galectins that regulate both immunity and placental development (Supplementary Table 4). To construct their gene phylogenies, we retrieved orthologous protein sequences of all seven human genes across twenty-one placental mammals and five outgroup species from Ensembl (human genome assembly GRCh38.p12) (Supplementary Table 5). By default, Ensembl provided the longest available isoform for each sequence. This can be problematic when evaluating signatures of positive selection, because sequence alignments based on the longest available isoforms tend to have increased amounts of misaligned positions that can lead to false detection of positive selection (Villanueva-Canas et al. 2013). Isoforms of similar length generate alignments of better quality and increase the accuracy of downstream molecular evolutionary analyses. Thus, we examined sequences that differed in length from the longest human isoform reported by Ensembl. These sequences were replaced if (1) Ensembl listed another isoform with a length more similar to the human isoform, or if (2) the NCBI Gene database listed an isoform with a more similar length. The accession numbers of all final isoforms included in the analysis are listed in Supplementary Table 6.

Some species lacked annotated orthologs in the Ensembl database, presumably due to gene loss, so the final sequence files for each galectin varied in the total number of sequences (Supplementary Table 7). This was most common for placental cluster galectins, which have experienced dramatic episodes of gene loss and gain during the evolution of primates (Than et al. 2009). Some species had multiple co-orthologs reported for the same human gene; for example, Ensembl reported three horse genes as co-orthologs for human *LGALS9*. All co-orthologs were included in the final sequence files (Supplementary Table 6).

Several non-primate galectins, mostly in artiodactyl species, were reported as orthologs for all three placental cluster galectins. These genes were included in the phylogenetic analysis to evaluate the evolutionary relationship between primate placental galectins and their non-primate counterparts. *LGALS13, LGALS14*, and *LGALS16* sequence files were combined into one sequence file with the non-primate sequences. In rare cases, placental cluster orthologs reported by Ensembl included genes previously identified as galectin pseudogenes by Than *et al.* (2009). These genes were not included in the final sequence files (see Supplementary Table 8 for a detailed description of all these exceptions and the corresponding reasons for their exclusion).

Protein sequences were aligned using the multiple sequence alignment software, MAFFT, version 7 (Katoh and Standley 2013). We produced five protein alignments: four for each galectin of the ancient group separately and one for the placental galectins and their non-primate orthologs together. Corresponding codon-based alignments were obtained using the sequence alignment webserver, PAL2NAL, version 14 (Suyama et al. 2006). Gene trees were reconstructed for each codon alignment using RAxML, version 8.2.4 (Stamatakis 2014). All trees for the ancient galectins were rooted using the sequence from the frog, *Xenopus tropicalis*, as the outgroup. The placental cluster gene tree was rooted with the opossum, *Monodelphis domestica*, as the outgroup.

We examined whether species trees were congruent with the gene sequence trees. Species trees were obtained using the Common Tree tool from the NCBI Taxonomy Database. The species trees contained two polytomies resulting from ambiguities in the placement of horse and green monkey on the mammalian phylogeny. We resolved these polytomies into bipartitions by enforcing horse to be a sister group to a clade containing pig, cow, sheep, and goat and by enforcing green monkey to be a sister group to a clade containing olive baboon and Macaques, in accordance with Perelman *et al*. (2011) and Shen *et al*. (2016). We applied the approximately unbiased (AU) test to evaluate whether the gene trees differed significantly from the corresponding species trees (Shimodaira 2002). Only the *LGALS8* gene tree agreed with its species tree (Supplementary Table 9). Thus, the *LGALS8* species tree was tested during the evolutionary rate analysis, while the gene tree was tested for all other galectins (Supplementary Figure 3).

### Evolutionary Rate Analysis

To estimate galectin nucleotide substitution rates across different timeframes in the evolution of placental mammals, we employed the codeML program from the phylogenetic analysis package, PAML, version 4.9c (Yang 2007). Codon frequency, transition/transversion rate ratio, and the nonsynonymous/synonymous rate ratio ω were estimated by codeML in all tests. ω is also annotated as *d*_*N*_*/d*_*S*_ and was used to estimate the strength of selection. Classic branch tests were used to evaluate different evolutionary scenarios in which ω varied between the placental stem branch, the clade of placental mammals, the clade of primates, and non-placental outgroup species (Supplementary Figure 4A). For tests involving the placental cluster galectins, branch tests were also used to evaluate scenarios in which ω varied between the primate stem branch, the non-primate outgroup clade, and the clades containing *LGALS13, LGALS14*, and *LGALS16* (Supplementary Figure 4B). Following Than *et al*. (2009), we assigned the placental stem branch the same ω as the outgroup species when not assigning it a unique ω. Similarly, we assigned the primate stem branch the same ω as the outgroup genes when not assigning it a unique ω. Six hypotheses were tested for each of the ancient galectin genes, and eight hypotheses were tested for the placental cluster galectin genes (Table 1). We used likelihood ratio tests to evaluate significance of each of the different evolutionary scenarios.

### Genotype Data for Population Genetics Analyses

To examine human galectin genes for signatures of recent positive selection and population differentiation, we retrieved human gene variant data from the 1000 Genomes Project Phase 3 release (1000 Genomes Project Consortium, 2015). Identifiers and genomic coordinates were downloaded for all seven human galectin genes from Ensembl (human genome assembly GRCh37.p13) via the python module PyEnsembl (http://www.hammerlab.org/). These coordinates were then used to extract corresponding genotype data for 1,668 individuals from the 1000 Genomes Project. These individuals comprised three different continental populations: African (661 individuals), European (503 individuals), and East Asian (504 individuals). We analyzed these three populations specifically because demographic parameters have been previously estimated for each of them (Gravel et al. 2011), thereby enabling the simulations of neutral evolution discussed in subsequent sections. We retrieved these gene variant data using a combination of the data retrieval tool, Tabix, version 0.2.6 (Li 2011), and a custom python script. We removed indels from these genotype data and excluded them from all subsequent analyses.

### Detection of Recent Positive Selection

We calculated three different metrics (Tajima’s *D, F*_ST_, and *H12*) to evaluate different signatures of positive selection across different time windows during recent human evolution. All three metrics were calculated for the genic region—defined by the genomic coordinates retrieved by PyEnsembl—of all seven galectin genes. Using the R package *PopGenome*, version 2.1.6 (Pfeifer et al. 2014), we calculated Tajima’s *D* to evaluate signatures left from selective events that occurred within the last ∼250,000 years (Tajima 1989; Sabeti et al. 2006). We then calculated weighted Weir & Cockerham’s fixation index (*F*_ST_) to detect signatures of population differentiation that occurred within the last ∼75,000 years (Weir and Cockerham 1984). Using the program *VCFtools* (Danecek et al. 2011), we calculated *F*_ST_ both gene-wise (i.e., for entire genes) and SNP-wise (i.e., for individual SNPs). One global comparison (i.e., considering all three populations at once) and three pairwise population comparisons (i.e., Africa vs. East Asia, Africa vs. Europe, and East Asia vs. Europe) were used to calculate four different *F*_ST_ values for every galectin gene and SNP.

Metrics like Tajima’s *D* are based on allele frequency spectra and detect signals left by a specific mode of positive selection known as a hard selective sweep, in which a single beneficial haplotype rises to near or full fixation in a population (Barrerio and Quintana-Murci 2010). Hard selective sweeps reduce nucleotide diversity in the genomic region neighboring the selected variant, as linked variants sweep through a population in a phenomenon called genetic hitchhiking (Barreiro and Quintana-Murci 2010).

Alternatively, positive selection may simultaneously act on *multiple* beneficial haplotypes through a soft selective sweep, which can bring several haplotypes to high frequency in a population (Pennings and Hermission 2006; Messer and Petrov 2013; Garud et al. 2015). Soft selective sweeps do not reduce nucleotide diversity and are better detected by metrics like the *H12* statistic, an index that considers the frequency of the two most common haplotypes in a population (Garud et al. 2015). Following Moon *et al*. (2018), we used a modified version of the original python script written by Garud and colleagues (Garud et al. 2015) to calculate a gene-wise *H12* for all seven analyzed galectin genes.

### Construction of Null Distributions

Demographic events and stochastic processes, such as genetic drift, can shape patterns of genetic variation and generate signatures similar to those of selection (Vitti et al. 2013), influencing the performance of each of the three metrics. To determine whether galectin gene variation can be accounted for by positive selection—as opposed to neutral evolution—we simulated null distributions for each metric based on a neutral model of human evolution. We compared observed galectin values against these null distributions to compute *p*-values. Galectins with a *p*-value lower than 0.05 (i.e., greater than the 5^th^ percentile of simulated values) for a given statistic were considered significant and likely to have experienced positive selection.

To construct null distributions, we first simulated neutrally evolved gene sequences with *SLiM*, version 2.4.1 (Haller and Messer 2017). Following Moon *et al*. (2018), we accounted for past demographic events by employing *SLiM*’s implementation of human demographic parameters previously calculated by Gravel *et al*. (2011) for Africans, East Asians, and Europeans. This simulation model first designates that the ancestral African population expanded from a size of 7,310 to 14,474 approximately 148,000 years ago. It further designates that the out-of-Africa migration occurred approximately 51,000 years ago, which was followed by the Eurasian split approximately 23,000 years ago. A population bottleneck followed the Eurasian split and reduced the European population to 1,032 individuals, and European and East Asian populations experienced exponential growth in population size over the last 23,000 years. The exponential coefficient was set at 0.0038 for Europeans and at 0.0048 for East Asians. Finally, the model assumes fixed rates for both recombination (1.0 x 10^-8^ recombination events per base pair) and mutation (2.4 x 10^-8^ mutations per base pair). More detailed demographic parameters regarding migration rates were also set in accordance with Moon *et al*. (2018).

We simulated the genotypes of 661, 503, and 504 individuals to reflect the number of individuals analyzed from the African, European, and East Asian populations, respectively. For each of the seven galectin genes, we simulated 1,000 sequences of the same length and then used these length-matched simulated sequences to calculate Tajima’s *D*, Weir and Cockerham’s *F*_ST_, and *H12*. These metrics were calculated for simulated sequences exactly as described for galectin genes.

### Analysis of Galectin SNPs

Examination of gene-wise patterns of selection may miss positively selected SNPs if they reside in otherwise neutrally evolving regions; however, site frequency spectrum-based metrics like Tajima’s *D* and haplotype-based metrics like *H12* were not designed for analysis of individual SNPs. By contrast, *F*_ST_ compares variation in SNP allele frequencies between populations (Barreiro and Quintana-Murci 2010) and can thus be applied to individual SNPs. To identify individual SNPs that are highly differentiated between populations, we compared the *F*_ST_ values of all galectin SNPs to the distributions of *F*_ST_ values of simulated SNPs using a *p*-value of 0.05 as the significance threshold. To focus on SNPs for which evidence of important functional effects is more likely, we filtered the SNPs with significant *F*_ST_ values by retaining only the those previously cited in the literature. We created this list by searching the Ensembl BioMart database (assembly GRCh38.p12) to identify which SNPs were previously cited. We further filtered this list by only retaining SNPs with a *putative* association to disease (i.e., prior studies explicitly claimed an association or functional link to a disease or phenotype). We submitted missense variants from this final list to the PolyPhen-2 webserver for preliminary variant effect prediction (Adzhubei et al. 2010).

### Estimation of Allele Ages

To estimate the age of each allele for SNPs with significant or near significant *F*_ST_ values that have previously been associated with human complex genetic disease, we generated minimum allele ages using parsimony ancestral reconstruction across the mammalian phylogenetic tree. We obtained sequence data for 30 mammals for each SNP from the Multiz Alignment of 30 mammals track in the UCSC genome browser using the UCSC Table Browser (Kent et al. 2002; Karolchik et al. 2004). We included additional allelic information from six great ape species using data from the Great Ape Project (Prado-Martinez et al. 2013). We further obtained ancient human sequencing information for six additional genomes from the Max Planck Institute for Evolutionary Anthropology (Meyer et al. 2012). A phylogenetic tree for all species was obtained from the TimeTree database (Kumar et al. 2017). We conducted parsimony ancestral state reconstruction for each allele using the phangorn package in R (Schliep 2011). The minimum age of each allele was assigned as the age of the first node ancestral to humans inferred not to contain the specified allele.

### Structural Analysis with RosettaCM

To predict the structural effects of missense variants, we used a comparative modeling approach with the RosettaCM application, which requires experimentally determined structures that can serve as templates. Comparative modeling uses the templates, which have a high degree of sequence similarity to the sequence of interest, to approximate the overall fold of the protein. Rounds of energetic minimization allow for minor conformational changes to accommodate the sequence differences and produce more accurate models (Bender et al. 2016). Currently available structures of most galectin proteins are incomplete, thereby limiting which variants we were able to successfully model. For example, Galectin-3’s N-terminal domain—containing both rs4644 and rs4652—lacks a published structure because it contains Pro-Gly repeats that make it highly flexible and difficult to crystallize (Ahmad et al. 2003). Of the four galectin candidates with variants passing previously discussed filters, Galectin-8 was found to be the most suitable protein for predicting variant effects. We thus employed RosettaCM from Rosetta version 3.8 to model the structural effects of a Galectin-8 variant associated with rheumatoid arthritis, along with three other tightly linked variants (Song et al. 2013).

We retrieved the PDB file for the template structure 4FQZ from the Protein Data Bank. 4FQZ was published in a study that structurally characterized Galectin-8 in complex with different common oligosaccharide ligands (Yoshida et al. 2012). 4FQZ represents a protease-resistant mutant form of Galectin-8 containing both the C-terminal and N-terminal CRDs. In this mutant form, the Galectin-8’s linker peptide was replaced by two amino acids (His-Met) to enable *in vivo* and *in vitro* experiments that are otherwise infeasible due to the linker’s susceptibility to proteolysis (Kim et al. 2013). Despite these mutations, this linker-shortened form serves as an excellent template because it represents a structure of Galectin-8 containing both CRDs. *In vivo* studies show that this mutated protein retains full activity and binds several of Galectin-8’s common binding partners (Yoshida et al. 2012; Kim et al. 2013), further supporting the use of 4FQZ as an appropriate template for comparative modeling.

We isolated Chain A from 4FQZ and retrieved its FASTA sequence from the PDB. The amino acid sequence of 4FQZ exactly matches the Galectin-8 reference sequence at all four variant positions, and thereby serves as the protein’s reference sequence. Two target sequences were aligned to this reference sequence: a sequence excluding the linker amino acids with four residue changes (F19>Y, R36>C, M56>V, and R158>S) to reflect the major Galectin-8 haplotype, and a sequence changing only F19>Y to reflect the hybrid haplotype. The latter change reflects the substitution caused by rs1126407, the variant usually studied in isolation. This experimental design allowed us to directly compare changes in the protein’s structure between the reference, hybrid, and major haplotypes.

We used Rosetta’s partial thread application to thread all target sequences over 4FQZ, providing an initial approximation of the protein structures for each target-template alignment by assigning coordinates from the template PDBs to the aligned residues in the target sequences. The threaded PDBs were then used as single inputs in separate iterations of the rest of the comparative modeling procedure. We used the RosettaCM application to model any missing residue densities, generating 10,000 all-atom models for each threaded PDB. All generated models were further minimized and refined with Rosetta’s dualspace relax protocol, bringing them closer to energetic minima. Parameters for both the initial comparative modeling and relaxation procedures are described in Supplementary File 1. We initially repeated the relaxation protocol on chain A from 4FQZ directly in order to compare it to the new models, but the resulting structure diverged dramatically from the best predicted model (RMSD ∼ 1.8). To resolve this issue, we subjected the reference sequence to the full procedure described above, generating and relaxing 10,000 hybridized structures based on the reference sequence.

After sorting models by overall energy (in Rosetta Energy Units), we selected the top 5,000 from each condition and clustered them with the clustering algorithm, Calibur (Li and Ng, 2010). The cutoff was adjusted until the largest cluster contained 20%-25% of the input models. We then selected the central model listed for the largest cluster as the best comparative model. These final models were compared to each other to evaluate structural differences between the reference haplotype, the major haplotype, and the hybrid haplotype. We used the PyMOL Molecular Graphics System, Version 2.1 Schrödinger, LLC., for all visual analyses, alignments, and RMSD calculations.

### Ensemble Analysis of Comparative Models

To account for structural variation among the top-scoring models of each condition and reflect the conformational flexibility of Galectin-8, we performed a clustering analysis of the top 100 scoring models from the reference, major, and hybrid haplotypes respectively. Using Calibur, we examined whether the top-scoring models of each haplotype grouped into different structure-based clusters. Model distances were represented by the RMSD in Å of Galectin-8’s N-terminal CRD spanning residues R45 to E89, the region containing eight residues known to directly interact with carbohydrate-bearing ligands (Yoshida et al. 2012). The cutoff was adjusted until approximately 50% of all tested models were assigned to a single cluster. We then aligned the central models of the two largest clusters from residues R45 to E89.

## Acknowledgements

Z.E. was supported by a Beckman Scholar Award. G.S. was supported by NIH grant T15LM007450. J.A.C. was supported by NIH grant R35GM127087 and the Burroughs Wellcome Fund Preterm Birth Initiative. A. R. was supported by the March of Dimes through the March of Dimes Prematurity Research Center Ohio Collaborative, the Burroughs Wellcome Fund Preterm Birth Initiative, and a Guggenheim fellowship. This work was conducted in part using the resources of the Advanced Computing Center for Research and Education at Vanderbilt University.

## Supplementary Material

**Supplementary Table 1:** Complete results from evolutionary analyses with codeML.

**Supplementary Table 2:** Highest calculated gene-wide *F*_*ST*_ values between human populations for galectins.

**Supplementary Table 3:** List of 138 SNPs whose *F*_*ST*_ values surpass significance threshold for positive selection.

**Supplementary Table 4:** Accession numbers of human galectin proteins used in the analysis.

**Supplementary Table 5:** Species included in the phylogenetic analysis.

**Supplementary Table 6:** Accession numbers of all protein sequences included in the phylogenetic analysis.

**Supplementary Table 7:** Number of sequences included in the evolutionary rate analysis for each galectin family member.

**Supplementary Table 8:** Exclusions/changes made during the identification of orthologs.

**Supplementary Table 9:** Results from the Approximately Unbiased test that compared galectin gene trees to the species phylogeny.

**Supplementary Figure 1:** Global alignment of the top-ten scoring models based on the major haplotype shows considerable structural variation.

**Supplementary Figure 2:** Models based on the major haplotype tend to occupy a different conformational space than models based on the reference haplotype or the hybrid haplotype.

**Supplementary Figure 3:** Gene trees used in codeML tests on *LGALS1, LGALS3*, and *LGALS9*. Bootstrap support values are included as branch labels.

**Supplementary Figure 4:** Phylogenies depicting the clades and branches tested in the evolutionary rate analysis for ancient galectins (A) and placental cluster galectins (B).

**Supplementary file 1:** Parameters used for comparative modeling with RosettaCM.

## References

Ahmad N, Gabius HJ, André S, Kaltner H, Sabesan S, Roy R, Liu B, Macaluso F, Brewer CF. 2003. Galectin-3 precipitates as a pentamer with synthetic multivalent carbohydrates and forms heterogeneous cross-linked complexes. J Biol Chem. 279:10841–10847.

Balan V, Nangia-Makker P, Schwartz AG, Jung YS, Tait L, Hogan V, Raz T, Wang Y, Yang ZQ, Wu GS, et al. 2008. Racial disparity in breast cancer and functional germ line mutation in galectin-3 (rs4644): a pilot study. Cancer Res. 68:10045–10050.

Banfer S, Schneider D, Dewes J, Strauss MT, Freibert SA, Heimerl T, Maier UG, Elsässer HP, Jungmann R, Jacob R. 2018. Molecular mechanism to recruit galectin-3 into multivesicular bodies for polarized exosomal secretion. Proc Natl Acad Sci USA. 115:E4396–E4405.

Barreiro L, Quintana-Murci L. 2010. From evolutionary genetics to human immunology: how selection shapes host defence genes. Nature Genet. 11:17–30.

de Boer R, Verweij N, van Veldhuisen DJ, Westra HJ, Bakker SJ, Gansevoort RT, Muller Kobold AC, van Gilst WH, Franke L, Mateo Leach I, van der Harst P. 2012. A Genome-Wide Association Study of Circulating Galectin-3. PLoS One. 7:e47385.

Chiariotti L, Salvatore P, Frunzio R, Bruni CB. 2002. Galectin Genes: Regulation of Expression. Glycoconj J. 19:441–449.

Cummings RD, Liu F. 2009. Galectins. In: Varki A, Cummings RD, Esko JD, et al., editors. Essentials of Glycobiology. Cold Spring Harbor (NY): Cold Spring Harbor Laboratory Press. Chapter 33.

Cummings RD, Liu F, Vasta GR. 2017. Galectins. In: Varki A, Cummings RD, Esko JD, et al., editors. Essentials of Glycobiology. Cold Spring Harbor (NY): Cold Spring Harbor Laboratory Press. Chapter 36.

Danecek P. 2011. The Variant Call Format and VCFtools. Bioinformatics. 27:2156–2158.

Di Lella S, Sundblad V, Cerliani JP, Guardia CM, Estrin DA, Vasta GR, Rabinovich GA. 2011. When Galectins Recognize Glycans: From Biochemistry to Physiology and Back Again. Biochemistry. 50:7842–7857.

Dumic J, Dabelic S, Flögel M. 2006. Galectin-3: an open-ended story. Biochim Biophys Acta. 4:616–635.

Fumagalli M, Sironi M, Pozzoli U, Ferrer-Admettla A, Pattini L, Nielsen R. 2011. Signatures of Environmental Genetic Adaptation Pinpoint Pathogens as the Main Selective Pressure through Human Evolution. PLoS Genet. 7:e1002355.

Garud NR. 2015. Recent Selective Sweeps in North American Drosophila melanogaster Show Signatures of Soft Sweeps. PLoS Genet. 11:e1005004.

Hamblin M, Thompson E, Di Rienzo, A. 2002. Complex Signatures of Natural Selection at the Duffy Blood Group Locus. Am J Hum Genet. 2:369–383.

Houzelstein D, Gonçalves IR, Fadden AJ, Sidhu SS, Cooper DN, Drickamer K, Leffler H, Poirier F. 2004. Phylogenetic Analysis of the Vertebrate Galectin Family. Mol Biol Evol. 21:117–1187.

Hu C, Chang SK, Wu CS, Tsai WI, Hsu PN. 2011. Galectin-3 gene (LGALS3) +292C allele is a genetic predisposition factor for rheumatoid arthritis in Taiwan. Int J Clin Rheumatol. 9:1227–1233.

Karolchik D, Hinrichs AS, Furey TS, Roskin KM, Sugnet CW, Haussler D, Kent WJ. 2004. The UCSC Table Browser data retrieval tool. Nucleic Acids Res. 32(Database issue):D493-6.

Katoh S, Standley D. 2013. MAFFT Multiple Sequence Alignment Software Version 7: Improvements in Performance and Usability. Mol Biol Evol. 30:772–780.

Kent WJ, Sugnet CW, Furey TS, Roskin KM, Pringle TH, Zahler AM, Haussler D. 2002. The human genome browser at UCSC. Genome Res. 12:996–1006.

Kim DS, Stanaway IB, Rajagopalan R, Bernbaum JC, Solot CB, Burnham N, Zackai EH, Clancy RR, Nicolson SC, Gerdes M, et al. 2012. Results of Genome-Wide Analyses on Neurodevelopmental Phenotypes at Four-Year Follow-Up following Cardiac Surgery in Infancy. PLoS One. 7:e45936.

Kim BW, Hong SB, Kim JH, Kwon DH, Song HK. 2013. Structural basis for recognition of autophagic receptor NDP52 by the sugar receptor galectin-8. Nature Commun. 4:1613–1613.

Kumar S, Stecher G, Suleski M, Hedges SB. 2017. TimeTree: A Resource for Timelines, Timetrees, and Divergence Times. Mol Biol Evol. 34:1812–1819.

Lewis SK, Farmer JL, Burghardt RC, Newton GR, Johnson GA, Adelson DL, Bazer FW, Spencer TE. 2007. Galectin 15 (LGALS15): A Gene Uniquely Expressed in the Uteri of Sheep and Goats that Functions in Trophoblast Attachment. Biol Reprod. 77:1027–1036.

Li H. 2011. Tabix: fast retrieval of sequence features from generic TAB-delimited files. Bioinformatics. 27:718–719.

Li S. Ng Y. 2010. Calibur: a tool for clustering large numbers of protein decoys. Bioinformatics. 11:25.

Madrigal-Matute J, Lindholt JS, Fernandez-Garcia CE, Benito-Martin A, Burillo E, Zalba G, Beloqui O, Llamas-Granda P, Ortiz A, Egido J, et al. 2014. Galectin-3, a Biomarker Linking Oxidative Stress and Inflammation With the Clinical Outcomes of Patients With Atherothrombosis. J Am Heart Assoc. 3:e000785.

Messer PW, Petrov DA. 2013. Population genomics of rapid adaptation by soft selective sweeps. Trends Ecol Evol. 28:10.

Meyer M, Kircher M, Gansauge MT, Li H, Racimo F, Mallick S, Schraiber JG, Jay F, Prufer K, de Filippo C, Sudmant PH, et al. 2012. A High-Coverage Genome Sequence from an Archaic Denisovan Individual. Science. 338:222–226.

Oakley M, Majam V, Mahajan B, Gerald N, Anantharaman V, Ward JM, Faucette LJ, McCutchan TF, Zheng H, Terabe M. 2012. Pathogenic Roles of CD14, Galectin-3, and OX40 during Experimental Cerebral Malaria in Mice. PLoS One. 4:e6793.

Ortiz JF, Rokas A. 2016. CTDGFinder: A Novel Homology-Based Algorithm for Identifying Closely Spaced Clusters of Tandemly Duplicated Genes. Mol Biol Evol. 34:215–229.

Pal Z, Antal P, Srivastava SK, Hullám G, Semsei AF, Gál J, Svébis M, Soós G, Szalai C, André S, et al. 2012. Non-synonymous single nucleotide polymorphisms in genes for immunoregulatory galectins: Association of galectin-8 (F19Y) occurrence with autoimmune diseases in a Caucasian population. Biochim Biophys Acta. 10:1512–1518.

Pennings PS, Hermisson J. 2006. Soft sweeps II--molecular population genetics of adaptation from recurrent mutation or migration. Mol Biol Evol. 23:1076–84.

Pennings PS, Hermisson J. 2006. Soft Sweeps III: The Signature of Positive Selection from Recurrent Mutation. PLoS Genet. 2:e186.

Perelman P, Johnson WE, Roos C, Seuánez HN, Horvath JE, Moreira MA, Kessing B, Pontius J, Roelke M, Rumpler Y, et al. 2011. A molecular phylogeny of living primates. PLoS Genet. 7:e1001342.

Pond S, Murrell B, Fourment M, Frost SD, Delport W, Scheffler K. 2011. A Random Effects Branch-Site Model for Detecting Episodic Diversifying Selection. Mol Biol Evol. 11:3033–3043.

Prado-Martinez J, Sudmant PH, Kidd JM, Li H, Kelley JL, Lorente-Galdos B, Veeramah K, Woerner A, O’Connor TD, Santpere G, et al. 2013. Great ape genome diversity. Nature. 499:471–475.

PyEnsembl. http://www.hammerlab.org/2015/02/04/exploring-the-genome-with-ensembl-and-python/.

Quinlan A, Hall I. 2010. BEDTools: a flexible suite of utilities for comparing genomic features. Bioinformatics. 26:841–842.

Rabinovich G, Toscano M. 2009. Turning ‘sweet’ on immunity: galectin–glycan interactions in immune tolerance and inflammation. Nature Immunol. 9:338–352.

Raj T, Kuchroo M, Replogle JM, Raychaudhuri S, Stranger BE, De Jager PL. 2013. Common risk alleles for inflammatory diseases are targets of recent positive selection. Am J Hum Genet. 92:517–529.

Ramos PS, Shaftman SR, Ward RC, Langefeld CD. 2014. Genes Associated with SLE Are Targets of Recent Positive Selection. Autoimmune Dis. doi:10.1155/2014/203435.

Ramos PS, Shedlock AM, Langefeld CD. 2015. Genetics of autoimmune diseases: insights from population genetics. J Hum Genet. 60:657–664.

Ruiz F, Scholz BA, Buzamet E, Kopitz J, André S, Menéndez M, Romero A, Solís D, Gabius HJ. 2014. Natural single amino acid polymorphism (F19Y) in human galectin-8: detection of structural alterations and increased growth-regulatory activity on tumor cells. FEBS J. 5:1446–1464.

Sabeti PC, Schaffner SF, Fry B, Lohmueller J, Varilly P, Shamovsky O, Palma A, Mikkelsen TS, Altshuler D, Lander ES. 2006. Positive natural selection in the human lineage. Science. 312: 1614–20.

Schliep, KP. 2011. phangorn: phylogenetic analysis in R. Bioinformatics. 27:592.

Sebban L, Ronen D, Levartovsky D, Elkayam O, Caspi D, Aamar S, Amital H, Rubinow A, Golan I, Naor D, et al. 2007. The involvement of CD44 and its novel ligand Galectin-8 in apoptotic regulation of autoimmune inflammation. J Immunol. 179:1225–1235.

Shen X, Salichos L, Rokas A. 2016. A Genome-Scale Investigation of How Sequence, Function, and Tree-Based Gene Properties Influence Phylogenetic Inference. Genome Biol Evol. 8:2268–2280.

Shi Y, Lin X, Chen G, Yan J, Ying M, Zheng X. 2011. Galectin-3 rs4652 A>C polymorphism is associated with the risk of gastric carcinoma and P-glycoprotein expression level. Oncol Lett. 14:8144–8149.

Shimodaira H. 2002. An Approximately Unbiased Test of Phylogenetic Tree Selection. Syst Biol. 51: 492–508.

Stamatakis A. 2014. RAxML Version 8:A tool for Phylogenetic Analysis and Post-Analysis of Large Phylogenies. Bioinformatics. 30: 1312–3.

Suyama M, Torrents D, Bork P. 2006. PAL2NAL: robust conversion of protein sequence alignments into the corresponding codon alignments. Nucleic Acids Res. 34:W609–12.

Than NG, Romero R, Erez O, Weckle A, Tarca AL, Hotra J, Abbas A, Han YM, Kim SS, Kusanovic JP, et al. 2008. Emergence of hormonal and redox regulation of galectin-1 in placental mammals: Implication in maternal–fetal immune tolerance. Proc Natl Acad Sci USA. 105:15819–15824.

Than NG, Romero R, Goodman M, Weckle A, Xing J, Dong Z, Xu Y, Tarquini F, Szilagyi A, Gal P, Hou Z, et al. 2009. A primate subfamily of galectins expressed at the maternal-fetal interface that promote immune cell death. Proc Natl Acad Sci USA. 106:9731–9736.

Than NG, Romero R, Kim CJ, McGowen MR, Papp Z, Wildman DE. 2012. Galectins: guardians of eutherian pregnancy at the maternal-fetal interface. Trends in Endocrinol and Metab. 23:23–32.

Than NG, Romero R, Xu Y, Erez O, Xu Z, Bhatti G, Leavitt R, Chung TH, El-Azzamy H, LaJeunesse C, Wang B, et al. 2014. Evolutionary origins of the placental expression of chromosome 19 cluster galectins and their complex dysregulation in preeclampsia. Placenta. 35:855–865.

Than NG, Romero R, Balogh A, Karpati E, Mastrolia SA, Staretz-Chacham O, Hahn S, Erez O, Papp Z, Kim CJ. 2015. Galectins: Double-edged Swords in the Cross-roads of Pregnancy Complications and Female Reproductive Tract Inflammation and Neoplasia. J Pathol Transl Med. 49:181–208.

The 1000 Genomes Project Consortium. 2015. A global reference for human genetic variation. Nature. 526:68–74.

Vasta GR. 2012. Galectins as pattern recognition receptors: structure, function, and evolution. Adv Exp Med Biol. 946:21–36.

Villanueva-Canas JL, Laurie S, Albà MM. 2013. Improving Genome-Wide Scans of Positive Selection by Using Protein Isoforms of Similar Length. Genome Biol Evol. 5:457–467.

Vitti JJ, Grossman SR, Sabeti PC. 2013. Detecting natural selection in genomic data. Annu Rev Genet. 47:97–120.

Weir B, Cockerham C. 1984. Estimating F-Statistics for the Analysis of Population Structure. Evolution. 38:1358–1370.

Yang Z. 2007. PAML 4:phylogenetic analysis by maximum likelihood. Mol Biol Evol. 24:1586-1591.

See PAML manual at http://abacus.gene.ucl.ac.uk/software/paml.html

Yoshida H, Yamashita S, Teraoka M, Itoh A, Nakakita S, Nishi N, Kamitori S. 2012. X-ray structure of a protease-resistant mutant form of human galectin-8 with two carbohydrate recognition domains. FEBS J. 279:3937–3951.

Zhang S, Moussodia RO, Vértesy S, André S, Klein ML, Gabius HJ, Percec V. 2015. Unraveling functional significance of natural variations of a human galectin by glycodendrimersomes with programmable glycan surface. Proc Natl Acad Sci USA. 18:5585–5590.

