## Supplementary Tables, Figures and Files for "The impact of natural selection on the evolution and function of placentally expressed galectins"

**Supplementary Table 1:** Complete results from evolutionary analyses with codeML.

| Gene | Hypothesis | Likelihood (-lnL) | *p*-value | ω values | | | |
|---|---|---|---|---|---|---|---|
| | | | | Outgroups | Placental Clade | Placental Stem | Primates |
| ***LGALS1*** | Null | 5353.76 | N/A | 0.15 | 0.15 | 0.15 | 0.15 |
| | 2 | 5344.95 | 1.5E-04 | 0.20 | 0.16 | 0.01 | 0.16 |
| | 3 | 5353.71 | 0.7584 | 0.15 | 0.16 | 0.15 | 0.16 |
| | 4 | 5345.53 | 5.0E-05 | 0.17 | 0.17 | 0.01 | 0.17 |
| | 5 | 5353.56 | 0.5284 | 0.15 | 0.15 | 0.15 | 0.18 |
| | 6 | 5353.55 | 0.8129 | 0.15 | 0.15 | 0.15 | 0.18 |
| ***LGALS3*** | Null | 7886.70 | N/A | 0.25 | 0.25 | 0.25 | 0.25 |
| | 2 | 7834.11 | 1.44E-23 | 0.07 | 0.39 | 0.21 | 0.39 |
| | 3 | 7834.84 | 2.33E-24 | 0.08 | 0.39 | 0.08 | 0.39 |
| | 4 | 7884.15 | 2.39E-02 | 0.26 | 0.26 | 0.08 | 0.26 |
| | 5 | 7873.09 | 1.82E-07 | 0.22 | 0.22 | 0.22 | 0.58 |
| | 6 | 7831.69 | 1.29E-24 | 0.07 | 0.35 | 0.07 | 0.57 |
| ***LGALS8*** | Null | 8161.84 | N/A | 0.19 | 0.19 | 0.19 | 0.19 |
| | 2 | 8161.09 | 0.469 | 0.17 | 0.20 | 0.27 | 0.20 |
| | 3 | 8161.66 | 0.5439 | 0.18 | 0.20 | 0.18 | 0.20 |
| | 4 | 8161.66 | 0.5506 | 0.19 | 0.19 | 0.24 | 0.19 |
| | 5 | 8153.47 | 4.29E-05 | 0.17 | 0.17 | 0.17 | 0.36 |
| | 6 | 8153.36 | 0.0002072 | 0.18 | 0.17 | 0.18 | 0.36 |
| ***LGALS9*** | Null | 14016.54 | N/A | 0.335 | 0.335 | 0.335 | 0.335 |
| | 2 | 14007.70 | 0.0001441 | 0.276 | 0.364 | 0.112 | 0.364 |
| | 3 | 14010.44 | 4.74E-04 | 0.235 | 0.364 | 0.235 | 0.364 |
| | 4 | 14009.79 | 2.36E-04 | 0.349 | 0.349 | 0.107 | 0.349 |
| | 5 | 14014.56 | 0.0461 | 0.318 | 0.318 | 0.318 | 0.396 |
| | 6 | 14009.85 | 0.001238 | 0.235 | 0.350 | 0.235 | 0.397 |

| | Hypothesis | Likelihood (-lnL) | *p*-value | Outgroups | Primate Stem | ***LGALS 13*** | ***LGALS 14*** | ***LGALS 16*** |
|---|---|---|---|---|---|---|---|---|
| **Placental Cluster Galectins** | Null | 4457.87 | N/A | 0.60 | 0.60 | 0.60 | 0.60 | 0.60 |
| | 2 | 4456.20 | 0.07 | 0.64 | 0.64 | 0.39 | 0.64 | 0.64 |
| | 3 | 4456.48 | 0.10 | 0.54 | 0.54 | 0.54 | 0.75 | 0.54 |
| | 4 | 4457.87 | 0.96 | 0.60 | 0.60 | 0.60 | 0.60 | 0.59 |
| | 5 | 4455.45 | 0.18 | 0.58 | 0.58 | 0.39 | 0.75 | 0.61 |
| | 6 | 4456.64 | 0.12 | 0.63 | 0.30 | 0.63 | 0.63 | 0.63 |
| | 7 | 4456.42 | 0.23 | 0.59 | 0.31 | 0.67 | 0.67 | 0.67 |
| | 8 | 4457.17 | 0.24 | 0.54 | 0.54 | 0.67 | 0.67 | 0.67 |

**Supplementary Table 2:** Highest calculated gene-wide $F_{ST}$ values between human populations for galectins.

| **Gene** | **Highest calculated $F_{ST}$** | **Corresponding Population Comparison** | ***p*-value** |
|---|---|---|---|
| *LGALS1* | 0.061 | African vs. European | 0.826 |
| *LGALS3* | 0.220 | African vs. East Asian | 0.193 |
| *LGALS8* | 0.114 | African vs. European | 0.508 |
| *LGALS9* | 0.056 | African vs. East Asian | 0.966 |
| *LGALS13* | 0.113 | European vs. East Asian | 0.385 |
| *LGALS14* | 0.207 | African vs. East Asian | 0.262 |
| *LGALS16* | 0.109 | African vs. East Asian | 0.637 |

**Supplementary Table 3:** List of 138 SNPs whose $F_{ST}$ values surpass significance threshold for positive selection.

| SNP | Highest calculated $F_{ST}$ |
|---|---|
| rs2853618 | 0.550327 |
| rs8012397 | 0.542395 |
| rs7160523 | 0.525929 |
| rs8004787 | 0.517382 |
| rs2075601 | 0.517382 |
| rs2075602 | 0.516312 |
| rs4652 | 0.515243 |
| rs1009978 | 0.460939 |
| rs1535502 | 0.449385 |
| rs2475830 | 0.440888 |
| rs1535503 | 0.417051 |
| rs1969746 | 0.413395 |
| rs1969745 | 0.413395 |
| rs2251642 | 0.404346 |
| rs7160110 | 0.381088 |
| rs72480733 | 0.373751 |
| rs6573004 | 0.360853 |
| rs2075598 | 0.359344 |
| rs11850516 | 0.359344 |
| rs3829283 | 0.355218 |
| rs1266382 | 0.354845 |
| rs6573006 | 0.35163 |
| rs6573005 | 0.35163 |
| rs1009977 | 0.349181 |
| rs2094102 | 0.346999 |
| rs4659682 | 0.341847 |
| rs4830 | 0.324832 |
| rs10755 | 0.308866 |
| rs911957 | 0.307188 |
| rs2758999 | 0.305125 |
| rs8698 | 0.304093 |
| rs2254487 | 0.303065 |
| rs72480725 | 0.299774 |
| rs2799410 | 0.298962 |
| rs10402453 | 0.296254 |
| rs16833769 | 0.288929 |
| rs2794787 | 0.28631 |
| rs74050921 | 0.283616 |
| rs16833828 | 0.281469 |
| rs1042275 | 0.278162 |
| rs56239009 | 0.265344 |
| rs819416 | 0.262326 |
| rs1041936 | 0.260833 |
| rs2853608 | 0.258246 |
| rs1266385 | 0.258239 |
| rs10424853 | 0.253716 |
| rs2799419 | 0.25261 |
| rs1266383 | 0.245405 |
| rs8011980 | 0.244786 |
| rs78733901 | 0.244056 |
| rs7159530 | 0.244056 |

| | |
|---|---|
| rs57196783 | 0.244056 |
| rs57064758 | 0.244056 |
| rs56358335 | 0.244056 |
| rs55848305 | 0.244056 |
| rs17128230 | 0.244056 |
| rs10498474 | 0.244056 |
| rs12077662 | 0.24257 |
| rs2794788 | 0.239896 |
| rs2794789 | 0.238116 |
| rs2758993 | 0.238116 |
| rs819419 | 0.235238 |
| rs74050920 | 0.231665 |
| rs75585784 | 0.227667 |
| rs56115130 | 0.226919 |
| rs78909201 | 0.225741 |
| rs3094268 | 0.221761 |
| rs10423105 | 0.221588 |
| rs10802546 | 0.215182 |
| rs67395212 | 0.21331 |
| rs28631884 | 0.205033 |
| rs10489790 | 0.204801 |
| rs10403702 | 0.203569 |
| rs3763959 | 0.200772 |
| rs361498 | 0.199185 |
| rs78813689 | 0.198457 |
| rs75328932 | 0.198457 |
| rs41305947 | 0.198457 |
| rs12080766 | 0.197843 |
| rs79344329 | 0.197752 |
| rs34759944 | 0.196113 |
| rs2853621 | 0.191786 |
| rs79484276 | 0.182802 |
| rs3820564 | 0.182668 |
| rs1266386 | 0.182346 |
| rs10418762 | 0.181255 |
| rs12092255 | 0.177426 |
| rs8109921 | 0.1736 |
| rs3922304 | 0.1736 |
| rs819428 | 0.173222 |
| rs112790242 | 0.171129 |
| rs7518541 | 0.16991 |
| rs538563061 | 0.168546 |
| rs11558455 | 0.166939 |
| rs10925158 | 0.165808 |
| rs56272109 | 0.165617 |
| rs7157768 | 0.165353 |
| rs28445525 | 0.164902 |
| rs2233708 | 0.164902 |
| rs2233706 | 0.164902 |
| rs10426654 | 0.164902 |
| rs7245574 | 0.164633 |
| rs2158964 | 0.164187 |
| rs4802053 | 0.162645 |

| | |
|---|---|
| rs2190913 | 0.161802 |
| rs2190912 | 0.161802 |
| rs7552576 | 0.160622 |
| rs73113014 | 0.159785 |
| rs75490731 | 0.15543 |
| rs73113027 | 0.154144 |
| rs113450127 | 0.154062 |
| rs12744425 | 0.15376 |
| rs77222245 | 0.153211 |
| rs2298098 | 0.151857 |
| rs16833850 | 0.148647 |
| rs142243957 | 0.146061 |
| rs76792063 | 0.14465 |
| rs75547343 | 0.14465 |
| rs35541195 | 0.144369 |
| rs2489151 | 0.143945 |
| rs4659681 | 0.143094 |
| rs2472126 | 0.141201 |
| rs1860133 | 0.141014 |
| rs73555858 | 0.140319 |
| rs56740824 | 0.139044 |
| rs1041939 | 0.138605 |
| rs10077 | 0.138605 |
| rs58158181 | 0.138599 |
| rs2273865 | 0.137905 |
| rs2273863 | 0.137905 |
| rs12078074 | 0.136964 |
| rs17753447 | 0.1367 |
| rs16833777 | 0.133011 |
| rs3094271 | 0.132156 |
| rs2233709 | 0.130876 |
| rs79501547 | 0.126621 |
| rs117424385 | 0.125013 |
| rs117212419 | 0.124019 |

**Supplementary Table 4:** Accession numbers of human galectin proteins used in the analysis.

| Protein Name | Accession Number |
|---|---|
| Galectin-1 | ENSP00000215909 |
| Galectin-3 | ENSP00000254301 |
| Galectin-8 | ENSP00000435460 |
| Galectin-9 | ENSP00000378856 |
| Galectin-13 | ENSP00000221797 |
| Galectin-14 | ENSP00000353893 |
| Galectin-16 | ENSP00000375904 |

**Supplementary Table 5:** Species included in the phylogenetic analysis.

| Common Name [a,b] | Scientific Name |
|---|---|
| *Human* | *Homo sapiens* |
| *Sumatran Orangutan* | *Pongo abelii* |
| European Rabbit | *Oryctolagus cuniculus* |
| **Duck-billed Platypus** | ***Ornithorhynchus anatinus*** |
| Domestic Goat | *Capra hircus* |
| Domestic Cow | *Bos taurus* |
| *Marmoset* | *Callithrix jacchus* |
| Domestic Dog | *Canis lupus familiaris* |
| *Green Monkey* | *Chlorocebus sabaeus* |
| Horse | *Equus caballus* |
| Domestic Cat | *Felis catus* |
| *Western Lowland Gorilla* | *Gorilla gorilla gorilla* |
| *Crab-eating Macaque* | *Macaca fascicularis* |
| *Rhesus Macaque* | *Macaca mulatta* |
| Prairie Vole | *Microtus ochrogaster* |
| **Gray Short-tailed Opossum** | ***Monodelphis domestica*** |
| Mouse | *Mus musculus* |
| Northern White-cheeked Gibbon | *Nomascus leucogenys* |
| Sheep | *Ovis aries* |
| *Chimpanzee* | *Pan troglodytes* |
| *Olive Baboon* | *Papio anubis* |
| Brown Rat | *Rattus norvegicus* |
| Wild Boar | *Sus scrofa* |
| **Red Junglefowl (chicken)** | ***Gallus gallus*** |
| **Green Anole** | ***Anolis carolinensis*** |
| **Western Clawed Frog** | ***Xenopus tropicalis*** |

[a] Italicized names in the left-hand column indicate the primate species included in the study of placental cluster galectins.
[b] Bolded names indicate non-placental outgroup species.

**Supplementary Table 6:** Accession numbers of all protein sequences included in the phylogenetic analysis.

| Gene Name | Species | Accession Number |
|---|---|---|
| *LGALS1* | Cow | ENSBTAP00000020080 |
| *LGALS1* | W.C. Frog | ENSXETP00000034893 |
| *LGALS1* | Opossum | ENSMODP00000000542 |
| *LGALS1* | Horse | ENSECAP00000005080 |
| *LGALS1* | Chicken | ENSGALP00000020275 |
| *LGALS1* | Green Monkey | ENSCSAP00000003561 |
| *LGALS1* | Dog | NP_001188417.1 |
| *LGALS1* | Rabbit | ENSOCUP00000019081 |
| *LGALS1* | Green Anole | ENSACAP00000015379 |
| *LGALS1* | Sheep | NP_001009287.1 |
| *LGALS1* | Platypus | ENSOANP00000013907 |
| *LGALS1* | Rhesus Macaque | NP_001162098.1 |
| *LGALS1* | Pig | NP_001001867.1 |
| *LGALS1* | Vole | ENSMOCP00000005985 |
| *LGALS1* | Rat | ENSRNOP00000013538 |
| *LGALS1* | C.E. Macaque | ENSMFAP00000031847 |
| *LGALS1* | Chimpanzee | ENSPTRP00000024707 |
| *LGALS1* | Gibbon | ENSNLEP00000030435 |
| *LGALS1* | Baboon | ENSPANP00000013280 |
| *LGALS1* | Cat | ENSFCAP00000007714 |
| *LGALS1* | Marmoset | XP_002743801.1 |
| *LGALS1* | Human | ENSP00000215909 |
| *LGALS1* | Gorilla | ENSGGOP00000011073 |
| *LGALS1* | Mouse | ENSMUSP00000086795 |
| *LGALS1* | Goat | ENSCHIP00000013044 |
| *LGALS1* | Orangutan | NP_001126310.1 |
| N/A (*LGALS1* co-ortholog) | C.E. Macaque | ENSMFAP00000013911.1 |
| N/A (*LGALS1* co-ortholog) | Dog | ENSCAFP00000041926.1 |
| N/A (*LGALS1* co-ortholog) | Rhesus Macaque | ENSMMUP00000054847.1 |
| N/A (*LGALS1* co-ortholog) | W.C. Frog | ENSXETP00000034889.3 |
| N/A (*LGALS1* co-ortholog) | W.C. Frog | ENSXETP00000034887.1 |
| N/A (*LGALS1* co-ortholog) | W.C. Frog | ENSXETP00000061154.1 |
| *LGALS3* | Opossum | ENSMODP00000014699 |
| *LGALS3* | W.C. Frog | NP_988986.1 |
| *LGALS3* | Cow | ENSBTAP00000041298 |
| *LGALS3* | Orangutan | XP_002824813.1 |
| *LGALS3* | Horse | ENSECAP00000004772 |
| *LGALS3* | Chicken | ENSGALP00000036506.2 |
| *LGALS3* | Dog | NP_001183972.1 |
| *LGALS3* | Green Monkey | ENSCSAP00000010782 |
| *LGALS3* | Green Anole | ENSACAP00000016002 |
| *LGALS3* | Rabbit | ENSOCUP00000002537 |
| *LGALS3* | Platypus | XP_001507412.1 |
| *LGALS3* | Sheep | ENSOARP00000022663 |
| *LGALS3* | Rhesus Macaque | ENSMMUP00000004746 |
| *LGALS3* | Rat | ENSRNOP00000070344 |
| *LGALS3* | Vole | ENSMOCP00000027347 |
| *LGALS3* | Pig | ENSSSCP00000030635 |
| *LGALS3* | Gibbon | ENSNLEP00000015544 |
| *LGALS3* | Chimpanzee | ENSPTRP00000010805 |
| *LGALS3* | C.E. Macaque | ENSMFAP00000006403 |
| *LGALS3* | Baboon | ENSPANP00000016951 |

| | | |
|---|---|---|
| *LGALS3* | Cat | ENSFCAP00000036138 |
| *LGALS3* | Mouse | ENSMUSP00000118169 |
| *LGALS3* | Goat | ENSCHIP00000008370 |
| *LGALS3* | Gorilla | XP_004055252.1 |
| *LGALS3* | Human | ENSP00000254301 |
| *LGALS3* | Marmoset | ENSCJAP00000058893 |
| *LGALS8* | Opossum | ENSMODP00000034917 |
| *LGALS8* | Orangutan | XP_002809312.1 |
| *LGALS8* | W.C. Frog | ENSXETP00000019750 |
| *LGALS8* | Cow | ENSBTAP00000018010 |
| *LGALS8* | Horse | ENSECAP00000013121 |
| *LGALS8* | Dog | ENSCAFP00000016207 |
| *LGALS8* | Green Monkey | XP_007988072.1 |
| *LGALS8* | Chicken | ENSGALP00000006727 |
| *LGALS8* | Rabbit | XP_017203199.1 |
| *LGALS8* | Sheep | XP_004021417.1 |
| *LGALS8* | Rhesus Macaque | ENSMMUP00000019833 |
| *LGALS8* | Vole | ENSMOCP00000025105 |
| *LGALS8* | Rat | ENSRNOP00000040412 |
| *LGALS8* | Pig | ENSSSCP00000057574 |
| *LGALS8* | Gibbon | ENSNLEP00000004047 |
| *LGALS8* | C.E. Macaque | ENSMFAP00000019740 |
| *LGALS8* | Baboon | ENSPANP00000010539 |
| *LGALS8* | Chimpanzee | ENSPTRP00000003612 |
| *LGALS8* | Cat | ENSFCAP00000028439 |
| *LGALS8* | Gorilla | ENSGGOP00000006759 |
| *LGALS8* | Goat | XP_017897728.1 |
| *LGALS8* | Mouse | ENSMUSP00000114200 |
| *LGALS8* | Marmoset | ENSCJAP00000032948 |
| *LGALS8* | Human | ENSP00000435460 |
| *LGALS9* | Opossum | ENSMODP00000024002.3 |
| *LGALS9* | W.C. Frog | ENSXETP00000062584 |
| *LGALS9* | Cow | ENSBTAP00000009025 |
| *LGALS9* | Orangutan | ENSPPYP00000009085 |
| *LGALS9* | Horse | XP_005597660.1 |
| *LGALS9* | Dog | ENSCAFP00000027504 |
| *LGALS9* | Green Monkey | ENSCSAP00000003779 |
| *LGALS9* | Rabbit | XP_017204586.1 |
| *LGALS9* | Green Anole | ENSACAP00000007994 |
| *LGALS9* | Sheep | ENSOARP00000017382 |
| *LGALS9* | Rhesus Macaque | ENSMMUP00000055016 |
| *LGALS9* | Rat | ENSRNOP00000017042 |
| *LGALS9* | Vole | XP_005349490.1 |
| *LGALS9* | Pig | ENSSSCP00000035946 |
| *LGALS9* | C.E. Macaque | ENSMFAP00000023855 |
| *LGALS9* | Gibbon | XP_004091505.1 |
| *LGALS9* | Baboon | ENSPANP00000019799 |
| *LGALS9* | Chimpanzee | ENSPTRP00000088151 |
| *LGALS9* | Cat | ENSFCAP00000008538 |
| *LGALS9* | Goat | XP_005693299.1 |
| *LGALS9* | Gorilla | ENSGGOP00000040205 |
| *LGALS9* | Mouse | ENSMUSP00000103904 |
| *LGALS9* | Marmoset | ENSCJAP00000022806 |

| | | |
|---|---|---|
| *LGALS9* | Human | ENSP00000378856 |
| N/A (*LGALS9* co-ortholog) | Green Monkey | ENSCSAP00000003021 |
| N/A (*LGALS9* co-ortholog) | Horse | ENSECAP00000020341 |
| N/A (*LGALS9* co-ortholog) | Horse | ENSECAP00000022279 |
| N/A (*LGALS9* co-ortholog) | Baboon | ENSPANP00000036288 |
| N/A (*LGALS9* co-ortholog) | Opossum | ENSMODP00000028818 |
| *LGALS13* | Green Monkey | ENSCSAP00000000165 |
| *LGALS13* | C.E. Macaque | ENSMFAP00000011778 |
| *LGALS13* | Chimpanzee | ENSPTRP00000018821 |
| *LGALS13* | Human | ENSP00000221797 |
| *LGALS13* | Gorilla | ENSGGOP00000020984 |
| *LGALS13* | Marmoset | ENSCJAP00000048794 |
| *LGALS13* | Rhesus Macaque | ENSMMUP00000004697 |
| *LGALS14* | Green Monkey | ENSCSAP00000000160 |
| *LGALS14* | Rhesus Macaque | ENSMMUP00000041406 |
| *LGALS14* | C.E. Macaque | ENSMFAP00000039363 |
| *LGALS14* | Baboon | ENSPANP00000046218.1 |
| *LGALS14* | Human | ENSP00000353893 |
| *LGALS14* | Gorilla | ENSGGOP00000004871 |
| *LGALS14* | Marmoset | ENSCJAP00000055940 |
| *LGALS16* | Green Monkey | ENSCSAP00000000170 |
| *LGALS16* | Rhesus Macaque | ENSMMUP00000006948 |
| *LGALS16* | C.E. Macaque | ENSMFAP00000012537 |
| *LGALS16* | Baboon | ENSPANP00000010742 |
| *LGALS16* | Chimpanzee | ENSPTRP00000043424 |
| *LGALS16* | Human | ENSP00000375904 |
| *LGALS16* | Gorilla | ENSGGOP00000022791.2 |
| *LGALS16* | Marmoset | ENSCJAP00000048794 |
| N/A (related to placental cluster galectins) | Opossum | ENSMODP00000017118 |
| N/A (related to placental cluster galectins) | Cow | ENSBTAP00000056054 |
| N/A (related to placental cluster galectins) | Cow | ENSBTAP00000020299 |
| *LGALS15* | Sheep | NP_001009238.1 |
| *LGALS13* (in Ensembl) | Pig | ENSSSCP00000003232 |
| *LGALS16* | Goat | ENSCHIP00000030006 |
| *LGALS15* | Goat | ENSCHIP00000023130 |
| N/A (related to placental cluster galectins) | Goat | ENSCHIP00000014979 |

**Supplementary Table 7:** Number of sequences included in the evolutionary rate analysis for each galectin family member.

| Protein name | Number of sequences | Species not included | Species with co-orthologs |
|---|---|---|---|
| Galectin-1 | 32 | N/A | C.E. Macaque, Dog, W.C. Frog, R. Macaque |
| Galectin-3 | 26 | N/A | N/A |
| Galectin-8 | 24 | Platypus, Green Anole | N/A |
| Galectin-9 | 29 | Chicken, Platypus | Green Monkey, Horse, Baboon, Opossum |
| Galectin-13 | 6 | Baboon, Orangutan, Marmoset | N/A |
| Galectin-14 | 7 | Chimpanzee, Orangutan | N/A |
| Galectin-16 | 7 | Marmoset, Orangutan | N/A |

**Supplementary Table 8:** Exclusions/changes made during the identification of orthologs.

| Protein, Species Name | Issue | Resolution |
|---|---|---|
| Galectin-1, Orangutan | Ensembl did not report an Orangutan ortholog for Galectin-1, though one is reported in Than et al. 2008 | Included the Orangutan Galectin-1 sequence from the NCBI Protein database |
| Galectin-13/16, Marmoset | Ensembl reported the same low-scoring ortholog[a] for both Galectin-13 and -16 for Marmoset, and Galectin-13 is not present in New World monkeys (Than et al. 2009). Relatively low sequence identity | Excluded ENSCJAP00000048794.2 from analysis |
| Galectin-13, Baboon | Low orthology confidence; gene only encodes truncated isoforms | Excluded ENSPANP00000032234.1 from analysis |
| Galectin-14, Orangutan | Low orthology confidence; Galectin-14 is reported as a pseudogene by Than et al. 2009 | Excluded ENSPPYG00000009975.1 from analysis |
| Galectin-16-like, C.E. Macaque | Two co-orthologs reported; one only encodes a severely truncated isoform | Excluded ENSMFAP00000024226.1 from analysis; retained other co-ortholog |
| Galectin-13/14/16 ortholog, Rabbit | Gene encodes bi-CRD protein twice the length of the prototype-class placental galectins | Excluded ENSOCUG00000027877 from analysis |

[a] Low-scoring orthologs are identified by a lack of high confidence in the Ensembl orthology reports.

**Supplementary Table 9:** Results from the Approximately Unbiased test that compared galectin gene trees to the species phylogeny.

| **Gene**[a] | ***p*-value** |
|---|---|
| *LGALS3* | 0.029 |
| *LGALS8* | 0.131 |

[a] *LGALS1* and *LGALS9* contained multiple co-orthologs in several species (e.g., three *LGALS9* co-orthologs in horse), so their gene trees were not comparable to species trees. Thus, *LGALS1* and *LGALS9* gene trees were used by default.

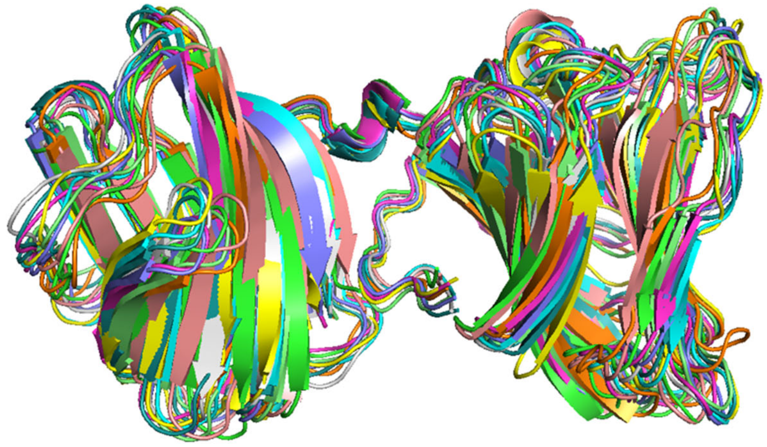

**Supplementary Figure 1:** Global alignment of the top-ten scoring models based on the major haplotype shows considerable structural variation.

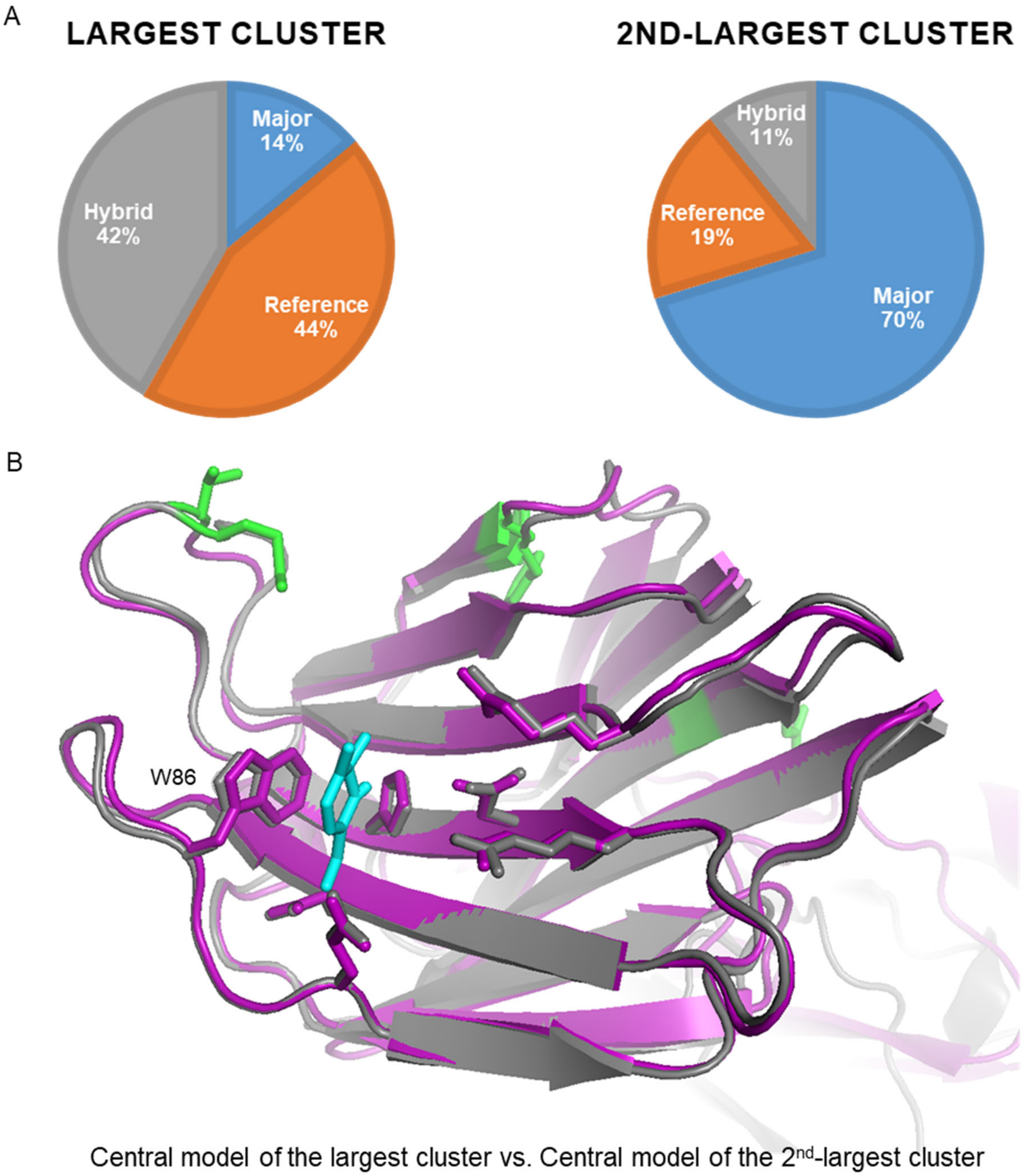


**Supplementary Figure 2:** Models based on the major haplotype tend to occupy a different conformational space than models based on the reference haplotype or the hybrid haplotype. (A) Model composition of the two largest clusters identified by Calibur: the largest cluster contains roughly equal numbers of models based on the reference haplotype and the hybrid haplotype, whereas models of the major haplotype occupy 70% of the 2nd-largest cluster. (B) Alignment of the N-terminal carbohydrate recognition domain (CRD) for the central models of the two largest clusters (RMSD=0.4 angstroms). The structure representing the 2$^{nd}$-largest cluster (of primarily major haplotype models) is colored purple. The structure representing the largest cluster (of primarily reference and hybrid haplotype models) is colored gray**.** The cyan-colored structure represents the approximate position typically occupied by carbohydrate ligands that bind the CRD.

**Supplementary Figure 3:** Gene trees used in codeML tests on *LGALS1*, *LGALS3*, and *LGALS9*. Bootstrap support values are included as branch labels.

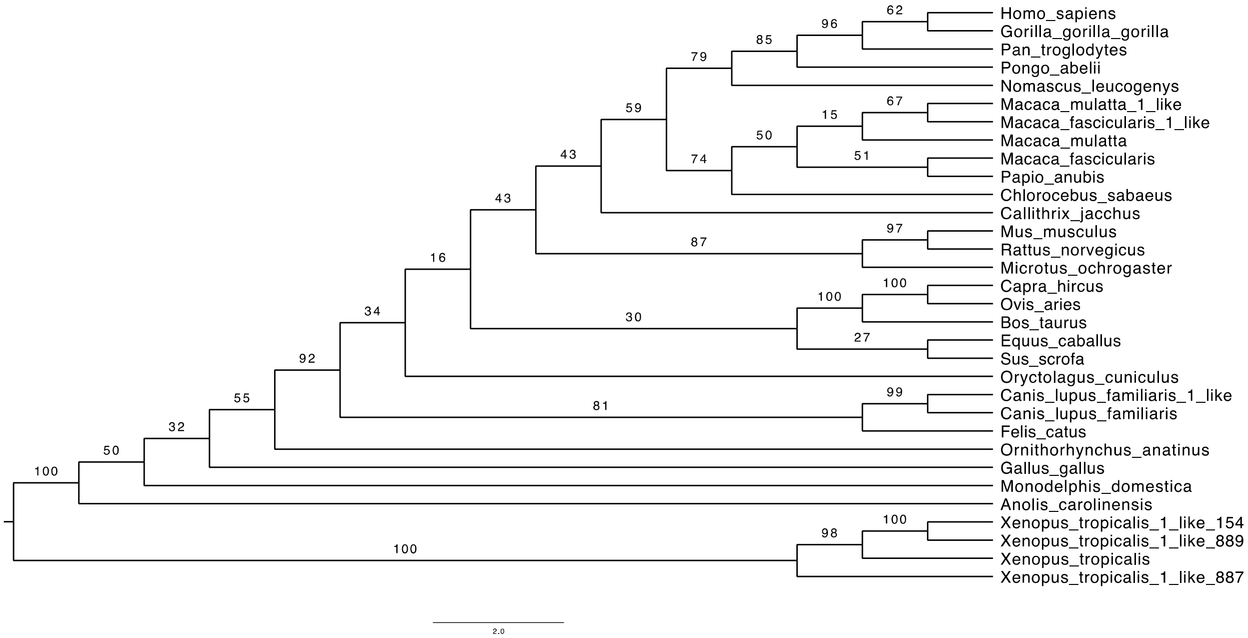


**Supplementary Figure 3A:** Gene tree used in codeML tests on *LGALS1*. The tree features several species containing multiple co-orthologs: *Xenopus tropicalis* (ENSXETP00000034889.3, ENSXETP00000034887.1, and ENSXETP00000061154.1), *Canis lupus familiaris* (ENSCAFP00000041926.1), *Macaca mulatta* (ENSMMUP00000054847.1), and *Macaca fascicularis* (ENSMMUP00000054847.1). Co-orthologs are annotated with the suffix "like" in the taxa labels.

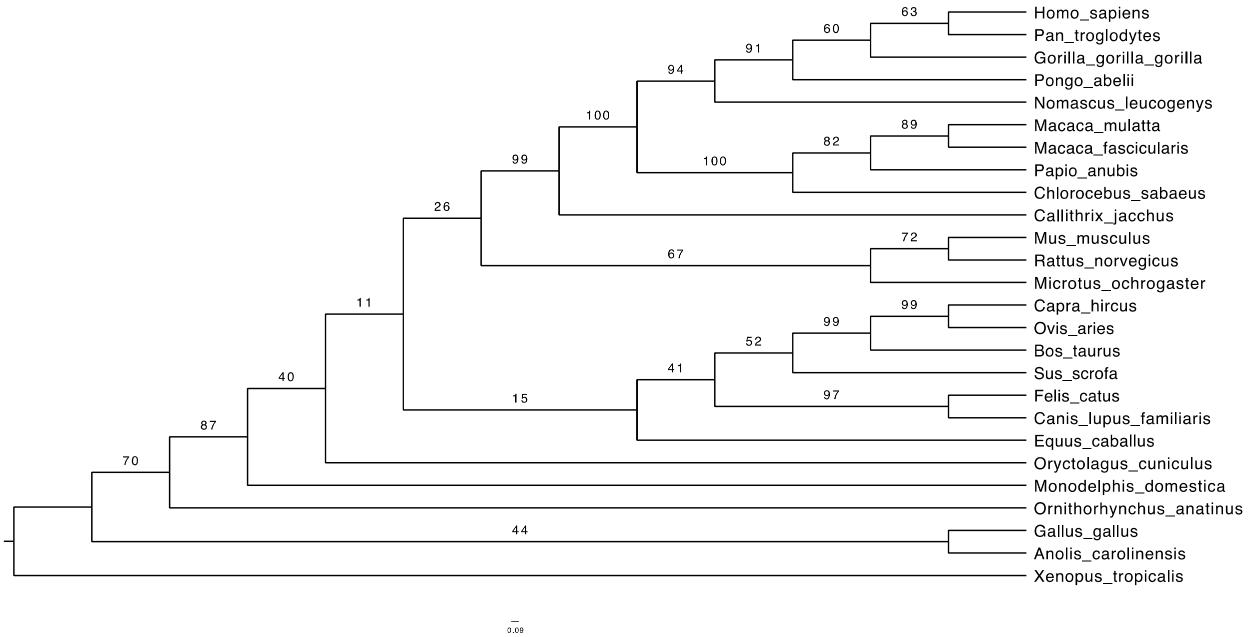


**Supplementary Figure 3B:** Gene tree used in codeML tests on *LGALS3*.

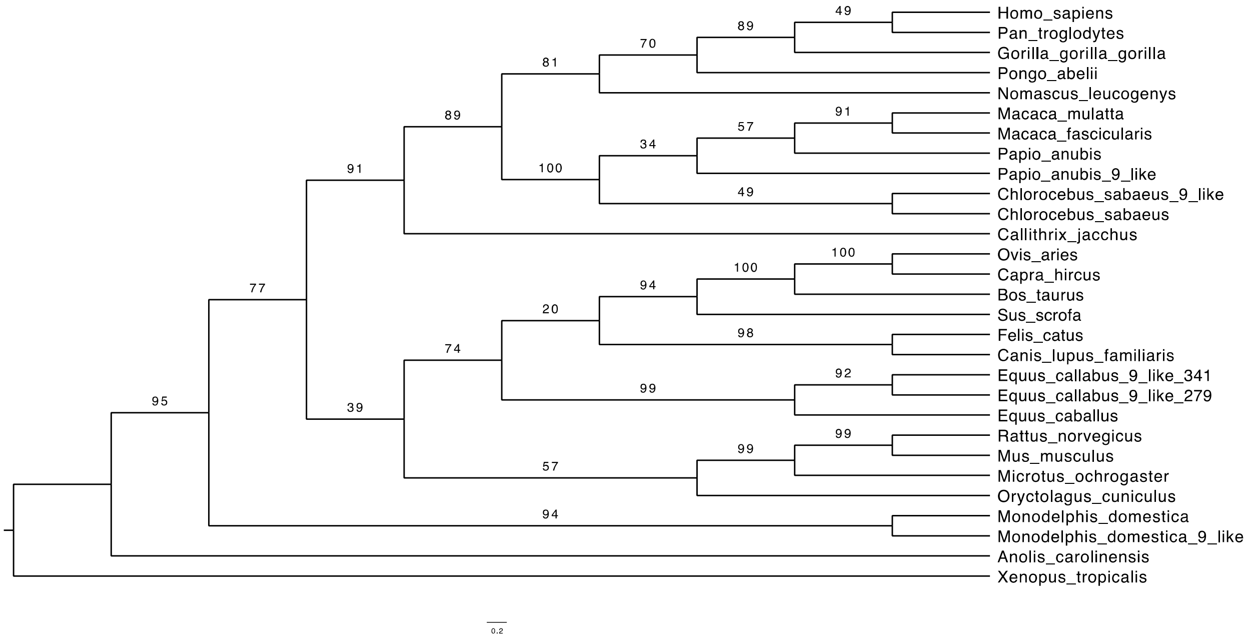


**Supplementary Figure 3C:** Gene tree used in codeML tests on *LGALS9*. The tree features several species containing multiple co-orthologs: *Papio anubis* (ENSPANP00000036288), *Chlorocebus sabaeus* (ENSCSAP00000003021), *Equus callabus* (ENSECAP00000020341 and ENSECAP00000022279), and *Monodelphis domestica* (ENSMODP00000028818). Co-orthologs are annotated with the suffix "like" in the taxa labels.

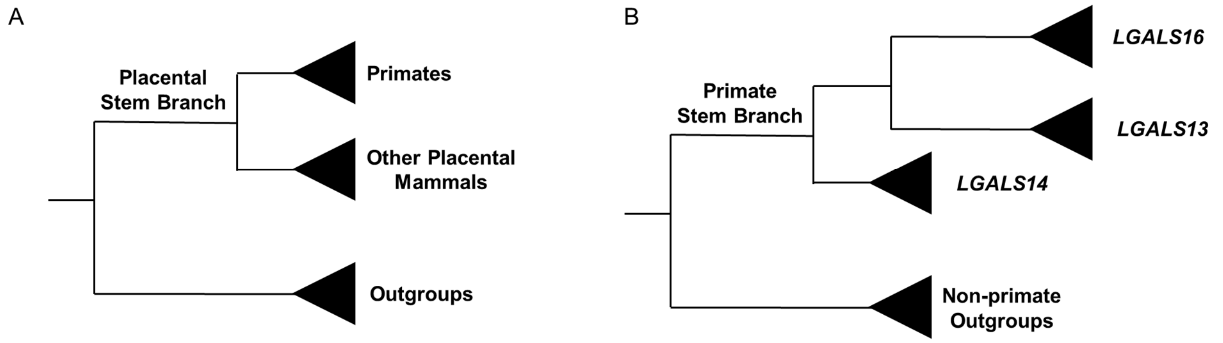


**Supplementary Figure 4:** Phylogenies depicting the clades and branches tested in the evolutionary rate analysis for ancient galectins (A) and placental cluster galectins (B).

**Supplementary file 1:** Parameters used for comparative modeling with RosettaCM.

The following command-line parameters were implemented during the hybridization step:
-nstruct 10000
-relax:minimize_bond_angles
-relax:min_type lbfgs_armijo_nonmonotone
-relax:jump_move true
-relax:default_repeats 2
-default_max_cycles 200

These parameters were used in conjunction with the Hybridize mover. The following score functions, weights, and parameters were incorporated into the mover:

-ScoreFunction name="stage1" weights="score3"
Reweight scoretype="atom_pair_constraint" weight="0.25"

-ScoreFunction name="stage2" weights="score4_smooth_cart"
Reweight scoretype="atom_pair_constraint" weight="0.25"

-ScoreFunction name="fullatom" weights="talaris2014_cart"
Reweight scoretype="atom_pair_constraint" weight="0.25"

-Hybridize name="hybridize" stage1_scorefxn="stage1" stage2_scorefxn="stage2" fa_scorefxn="fullatom" batch="1" stage1_increase_cycles="1.0" stage2_increase_cycles="1.0" linmin_only="0" realign_domains
Template pdb="4fqz_A_thread.pdb" weight="1.0" cst_file="AUTO"

Hybridization was followed by relaxation using Rosetta's dualspace relax protocol with these command-line parameters:

-relax:dualspace
-relax:minimize_bond_angles
-set_weights cart_bonded .5 pro_close 0
-default_max_cycles 200
-out:file:fullatom
-out:pdb
